# Identifying predictors of HPV-related head and neck squamous cell carcinoma progression and survival through patient-derived models

**DOI:** 10.1101/652537

**Authors:** ND Facompre, P Rajagopalan, V Sahu, AT Pearson, KT Montone, CD James, FO Gleber-Netto, GS Weinstein, J Jalaly, A Lin, AK Rustgi, H Nakagawa, JA Califano, CR Pickering, EA White, B Windle, IM Morgan, RB Cohen, PA Gimotty, D Basu

**Author notes:** Corresponding Author: Devraj Basu, 3400 Spruce Street, 5 Ravdin/Silverstein, Philadelphia, PA 19104.

## Abstract

Therapeutic innovation for human papilloma virus-related (HPV+) head and neck squamous cell carcinomas (HNSCCs) is impaired by inadequate biomarkers and preclinical models. This study addressed both limitations using the largest panel of HPV+ HNSCC patient-derived xenografts (PDXs) and organoids described to date. Whole exome profiles of the PDXs were compared to those of HPV+ human tumors and cell lines, and genetic features of the models were analyzed relative to their growth properties and outcomes of their patients of origin. PDX engraftment enriched for negatively prognostic NOTCH1 mutations while preserving multiple features lost in existing cell lines, including *PIK3CA* mutations, *TRAF3* deletion, and absence of *EGFR* amplification. Observation of more mutations in faster-growing models facilitated identification of an association between mutational burden and local progression in both HPV+ and HPV-HNSCCs. Reduced E7 and p16^INK4A^ levels found in a PDX from a lethal case led to detection of a similar profile among recurrent HPV+ HNSCCs in two patient cohorts, where low E2F target gene expression downstream of E7 predicted recurrence and mortality. Our findings bridge a critical gap in preclinical models for HPV+ HNSCCs and simultaneously reveal novel applications for mutational burden and E2F target dysregulation in biomarker development.

## INTRODUCTION

The human papilloma virus-related (HPV+) subtype of head and neck squamous cell carcinoma (HNSCC) occurs in a younger patient demographic than other HNSCCs and is rising in incidence (1). A major therapeutic objective for these patients is to decrease the severe treatment-related morbidities that persist after radiation and cytotoxic chemotherapy (2) while maintaining their generally favorable survival outcomes (3). However, attempting to reduce treatment toxicity for this disease through therapy de-escalation carries significant risk. For instance, treatment failures increased in two recent phase III trials evaluating therapeutic efficacy of cetuximab as an alternative to standard cisplatin (4, 5). Improving clinical trial design for this disease is presently impeded by the limited risk stratification offered by 8^th^ edition AJCC staging (6) and absence of molecular biomarkers. These factors are barriers both to identifying the optimal HPV+ HNSCC patients for therapy de-escalation and to developing distinct, more effective treatments for high risk cases. Among molecular prognostic features that could potentially guide treatment (7-10), *NOTCH1* loss of function is thus far the only mutation shown to predict poor outcomes in a published cohort of HPV+ HNSCCs (10).

A second obstacle to therapeutic innovation for HPV+ HNSCCs is the paucity of experimental models that accurately recapitulate their biology in humans. Poor growth of these tumors *ex vivo* delayed the creation of human cancer cell lines from them, and the cell lines that are now available lack multiple genetic traits that distinguish HPV+ HNSCCs from their HPV-counterparts (11). Artifacts in existing HPV+ cell lines include loss of the canonical *PIK3CA* activating mutations and *TRAF3* deletions that are more abundant in the HPV+ subtype of HNSCC. Most HPV+ HNSCC cell lines also acquire *EGFR* amplifications, which are uncommon in HPV+ HNSCCs, as well as copy gains in the 3q region spanning *TP63, SOX2*, and *PIK3CA* (11,12). Infrequent surgical treatment of HPV+ HNSCCs in the past and their intrinsically poor engraftment rates in immune-deficient mice (13) have impeded development of patient-derived xenograft (PDX) models. As a result, current preclinical models are mostly derived from cell lines that have limited ability to assess strategies to reduce treatment toxicity for low risk cases or evaluate intensified approaches for high risk cases.

The improved molecular fidelity of PDXs and organoids over cancer cell lines is well established (14,15) and justified our pursuit of such models despite potential pitfalls of generating them from HPV+ HNSCCs. In particular, the <25% stable engraftment rate observed by us (13) raised the possibility that any models that were created would also contain major molecular selection biases. Although such engraftment biases may prevent PDXs from capturing some tumor genotypes, PDXs can also enrich for other clinically relevant molecular subgroups, including those with more aggressive behavior (15-18). For instance, rapid growth of early stage HPV-HNSCCs upon engraftment to mice was recently shown to be predictive of recurrence and mortality for that disease subtype (19). These observations informed our intensive effort to generate a limited number of PDXs from HPV+ HNSCCs in order to provide preclinical models for aggressive tumor phenotypes and identify molecular traits that could aid risk stratification in the clinic.

This study aimed to characterize the largest panel of HPV+ HNSCC PDXs and organoids described to date and leverage these models to advance molecular risk stratification. Comparative analysis of whole exome sequence (WES) data between HPV+ and HPV-PDXs was used to evaluate the HPV+ models for retention of genetic traits specific to HPV+ HNSCCs, including those not captured by HPV+ cell lines. Cancer-related mutations in the PDXs were assessed for enrichment relative to their expected frequencies derived from the Cancer Genome Atlas (TCGA) to identify potentially aggressive genotypes that promote engraftment. A higher mutational burden observed in the HPV+ PDXs and organoids that grew most efficiently was further tested for ability to predict tumor progression and outcomes in patients. In addition, distinctive molecular features found in a PDX from a rapidly lethal HPV+ case served to identify a gene expression pattern with prognostic utility in two published HPV+ HNSCC cohorts.

## RESULTS

### PDXs retain genetic hallmarks of HPV+ HNSCC and avoid artifacts seen in HPV+ cell lines

Oropharyngeal primary HNSCCs or lymph node metastases that were positive for p16^INK4A^ by immunohistochemistry (IHC) were used to generate 9 stable HPV+ PDXs (Supplemental Table S1). Whole exome sequencing (WES) was completed for 8 HPV+ PDXs, and targeted sequencing was performed selectively on a 9^th^ HPV+ PDX that failed to provide full WES due to excess mouse DNA. Sequenced models all contained HPV-16 DNA and expressed HPV-16 transcripts by qPCR (Supplemental Figure S1). WES profiles of the HPV+ PDXs were compared to those of the 53 HPV+ oropharyngeal HNSCCs in TCGA (Supplemental Table S2)(20). Despite the small sample size, the PDXs captured mutations in 67% of the genes in the COSMIC Cancer Gene Census (CGC) that are mutated in HPV+ TCGA cases at a frequency of ≥10% (Figure 1A). WES simultaneously performed on 11 PDXs from HPV-HNSCCs (Supplemental Table S1) provided controls to help highlight distinguishing features of HPV+ HNSCCs preserved in the models. Genetic differences between HPV+ and HPV-HNSCCs previously reported in at least two studies (21-30) (Supplemental Table S3) were broadly preserved between the two groups of PDXs (Figure 1B). Distinguishing features of the HPV+ models included a paucity of *TP53* and *CDKN2A* alterations (21-26, 28, 29) and multiple KMT2C mutants (25,30). The HPV+ PDXs also avoided *EGFR* copy gains, which are rare in human HPV+ HNSCCs but are typically acquired by their cell lines, and captured a characteristic *TRAF3* deep deletion not found in HPV+ cell lines (11) (Figure 1C). Furthermore, the HPV+ PDXs retained the increase in alterations predicted to activate the PI3-kinase pathway that are a hallmark of HPV+ HNSCC (Figure 1D). *PIK3CA* hotspot mutations, which are lost in HPV+ cell lines (11,12), were identified in 3 HPV+ PDXs, including a canonical activating E545K helical domain mutant (Figure 1E, Supplemental Table S4). The two other mutants (E970K and N1044K) were in the catalytic domain and are previously described as hotspots in a population-scale cohort of tumor samples (31,32), with N1044K identified by targeted sequencing of the HPV+ PDX that failed WES. Finally, the 3q amplicon spanning *PIK3CA, TP63*, and *SOX2* that appears de novo during HNSCC cell line generation (11,12) was not enriched the HPV+ PDXs (Figure 1F). Together, these findings supported the superior fidelity of the PDXs over cell lines in representing key genetic traits of HPV+ HNSCC and thus their distinct utility as HPV+ cancer models.

**Figure 1.**
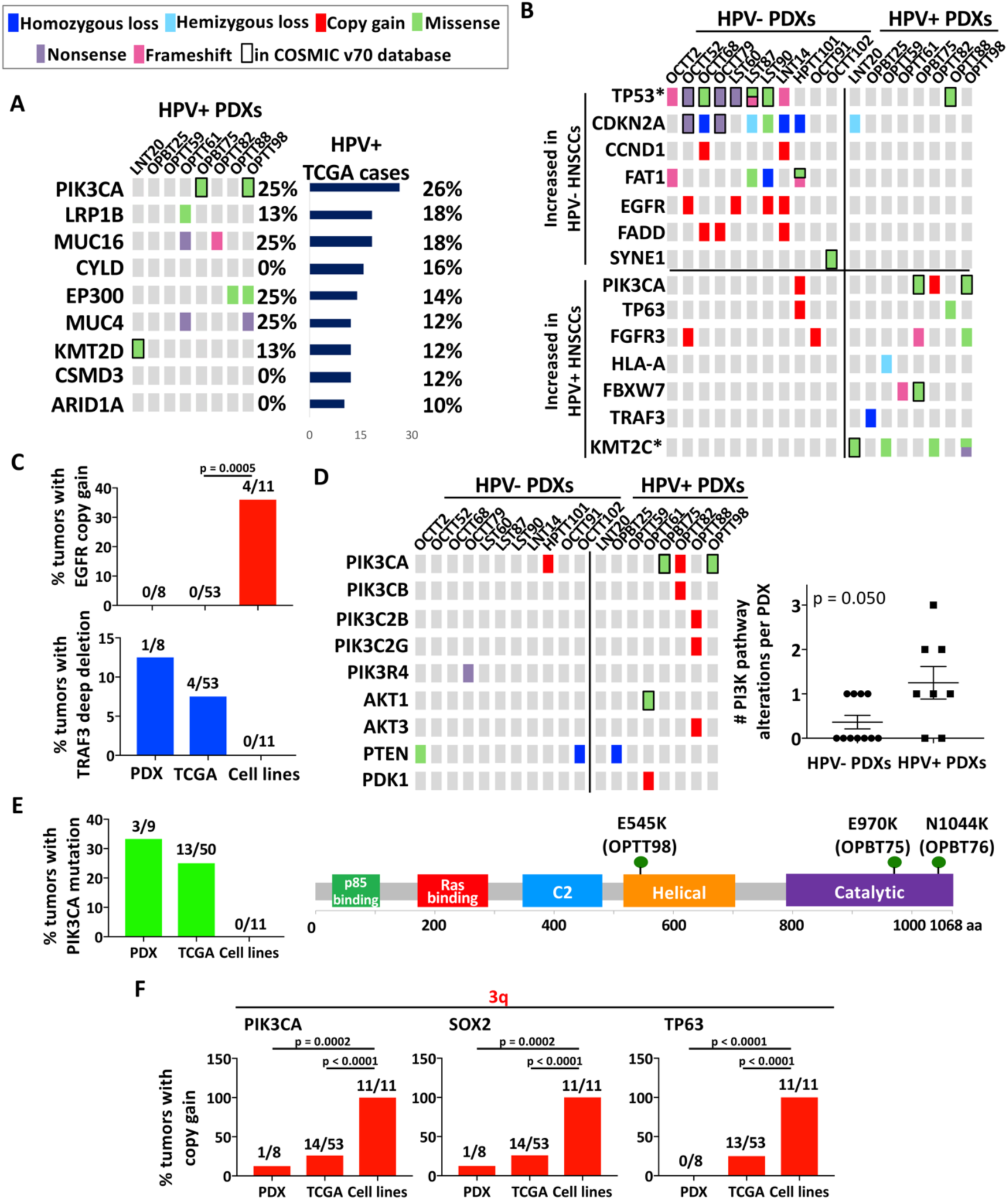
PDXs retain genetic hallmarks of HPV+ HNSCC and avoid artifacts seen in HPV+ cell lines. **A**. PDX profiles for the CGC genes mutated in ≥10% of HPV+ HNSCCs in TCGA. **B**. Frequencies in PDXs of mutations and CNAs that differ between HPV- and HPV+ HNSCCs. Unadjusted p-values determined by Fisher’s exact test. *p<0.025 **C**. Frequencies of EGFR and TRAF3 CNAs in HPV+ PDXs, TCGA cases, and cell lines. p-values determined by Fisher’s exact test. **D**. Mutations and CNAs in proximal components of the PI3K/Akt pathway compared between HPV- and HPV+ PDXs (left). Number of altered PI3K pathway genes per tumor in HPV-vs. HPV+ PDXs (right). p-value determined by two-tailed Student’s t-test assuming unequal variances. **E**. Frequencies of PIK3CA mutations in HPV+ PDXs, TCGA cases, and cell lines (left). Positions of *PIK3CA* mutations in HPV+ PDXs (right). **F**. Frequencies of 3q gene copy gains in HPV+ PDXs, TCGA cases, and cell lines. p-values determined by Fisher’s exact test.

### HPV+ PDXs capture a higher frequency of Notch pathway mutations than found in patients

Despite overall fidelity of HNSCC PDXs to the genetic landscape of HPV+ tumors, low engraftment rates in immunodeficient mice (13) predicted molecular selection biases in the PDXs. To evaluate for such effects that might provide models for more aggressive tumor subtypes, mutations in CGC genes that occurred in at least two HPV+ PDXs were compared in frequency to their counterparts in TCGA (Figure 2A). *KMT2C* mutations were increased in the PDXs relative to TCGA but are found at high frequency in other HPV+ HNSCC series (25,30). By contrast, significantly more *NOTCH1* mutants with predicted loss of function were found in the PDXs relative to TCGA and the three other large published cohorts (Figure 2B). The mutations occurred predominantly in the N-terminal EGF ligand-binding domain (Figure 2C), like those recently reported to be negatively prognostic (10). These mutations were accompanied by a significant increase in alterations across the other Notch receptors and ligands in aggregate (Figure 2D). A similar increase was not evident in the HPV-PDXs (Supplemental Figure S2), despite comparable background frequencies of Notch pathway mutations in HPV+ and HPV-HNSCCs (24). To evaluate whether the increase in HPV+ models arose through selective engraftment, targeted sequencing was used to detect the frequency of *NOTCH1* mutants in the population of HPV+ tumors from which engraftment was attempted. A trend toward more *NOTCH1* mutations in tumors that stably engrafted (Figure 2E) was accompanied by higher mutant *NOTCH1* allele frequencies in the established PDXs relative to their tumors of origin (Supplemental Table S5). For one HPV+ PDX (LNT20), the *NOTCH1* mutation was undetectable in the tumor of origin, indicating either strong clonal enrichment or *de novo* mutation during engraftment or initial passage. Taken together, these findings demonstrate that PDXs efficiently capture alterations predicted to inactivate the Notch pathway in HPV+ HNSCCs. A paucity of similar changes in HPV+ cell lines (11,12) establishes the PDXs as compelling models for this class of molecular alteration, which likely plays a major role in HPV+ HNSCC biology (33) and may have negative prognostic significance (10).

**Figure 2.**
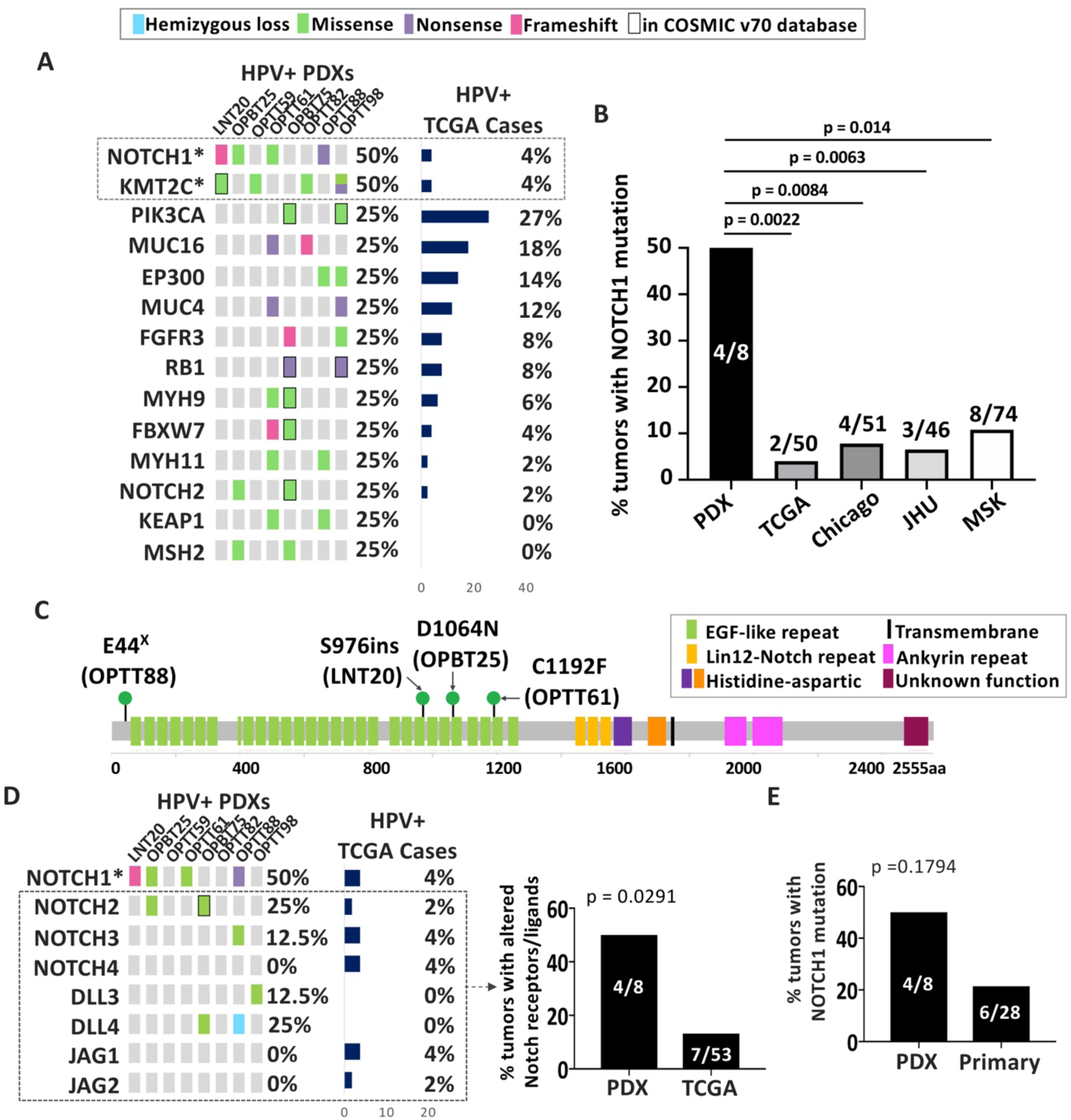
HPV+ PDXs capture a higher frequency of Notch pathway mutations than found in patients. **A**. Genes in CGC mutated in two or more HPV+ PDXs compared to the same genes in HPV+ TCGA HNSCCs. Unadjusted p-values: *p<0.005 **B**. Frequencies of NOTCH1 mutations in HPV+ PDXs compared with cases in the HPV+ TCGA, Chicago (25), JHU (30), and MSK (10) patient cohorts. p-values were determined by Fisher’s exact test. **C**. Positions of *NOTCH1* mutations in HPV+ PDXs. **D**. Mutations and CNAs in NOTCH receptors and ligands in HPV+ PDXs with predicted loss of function (left). Unadjusted p-values: *p<0.005 Frequency of mutations and/or CNAs in NOTCH receptor (excluding NOTCH1) or NOTCH ligand genes in HPV+ PDXs vs. TCGA (right). **E**. Percent of tumors with mutant NOTCH1 in HPV+ PDXs and HPV+ human HNSCCs from which engraftment was attempted. p-values were determined by Fisher’s exact test.

### High mutational burden is associated with efficient growth of HPV+ PDXs and organoids

To identify genetic features that might impact HPV+ HNSCC progression, growth properties of the HPV+ PDXs were evaluated for relationships with their mutational profiles. PDX growth curves (Figure 3A) were quantified by averaging the regression coefficients from two consecutive passages, and growth rates were expressed as slopes weighted by the variability among three or more replicates (Supplemental Table S6). The relationship between mutation profile and growth rate was evaluated using principal component analysis (PCA), and a relatedness dendrogram was created based on the Euclidian distances among allele-frequency weighted mutation vectors (Figure 3B, bottom). The dendrogram segregated fast, intermediate, and slow-growing PDXs (Figure 3B, top), whose PCA grouping is also illustrated on a projection plot (Supplemental Figure S3). Although the small PDX sample size was ill-suited to identifying growth associations with individual genes, a significant positive association was apparent between growth rate and the total number of called mutations, described here as tumor mutational burden (TMB) (Figure 3C). Furthermore, creation of *in vitro* organoid models from the PDXs revealed associations between organoid growth efficiency and both *in vivo* growth rate (Figure 3D, left) and TMB (right, Supplemental Figure S4). These *in vitro* proliferative features indicated the PDX growth rates to be tumor-autonomous features and not artifacts of the mouse milieu. Thus, the CGC gene list was used to interrogate whether mutations in the subset of these cancer genes account for the growth rate differences. Surprisingly, subtracting mutations in known oncogenes and tumor suppressors from the TMB count failed to diminish the strength of the growth association (Figure 3E). This finding was consistent with alterations outside the known cancer genome driving PDX growth or simply an accumulation of passenger mutations in rapidly growing models. The latter possibility was supported by the growth association also being present for point substitutions excluded by the mutation-calling pipeline, including missense mutations without damaging R-SVM scores (34) (Figure 3F, left) and synonymous mutations (right). Collectively, these findings revealed that faster growing HPV+ models harbor more passenger mutations and indicated potential utility for TMB as a predictor of tumor-autonomous HPV+ HNSCC progression.

**Figure 3.**
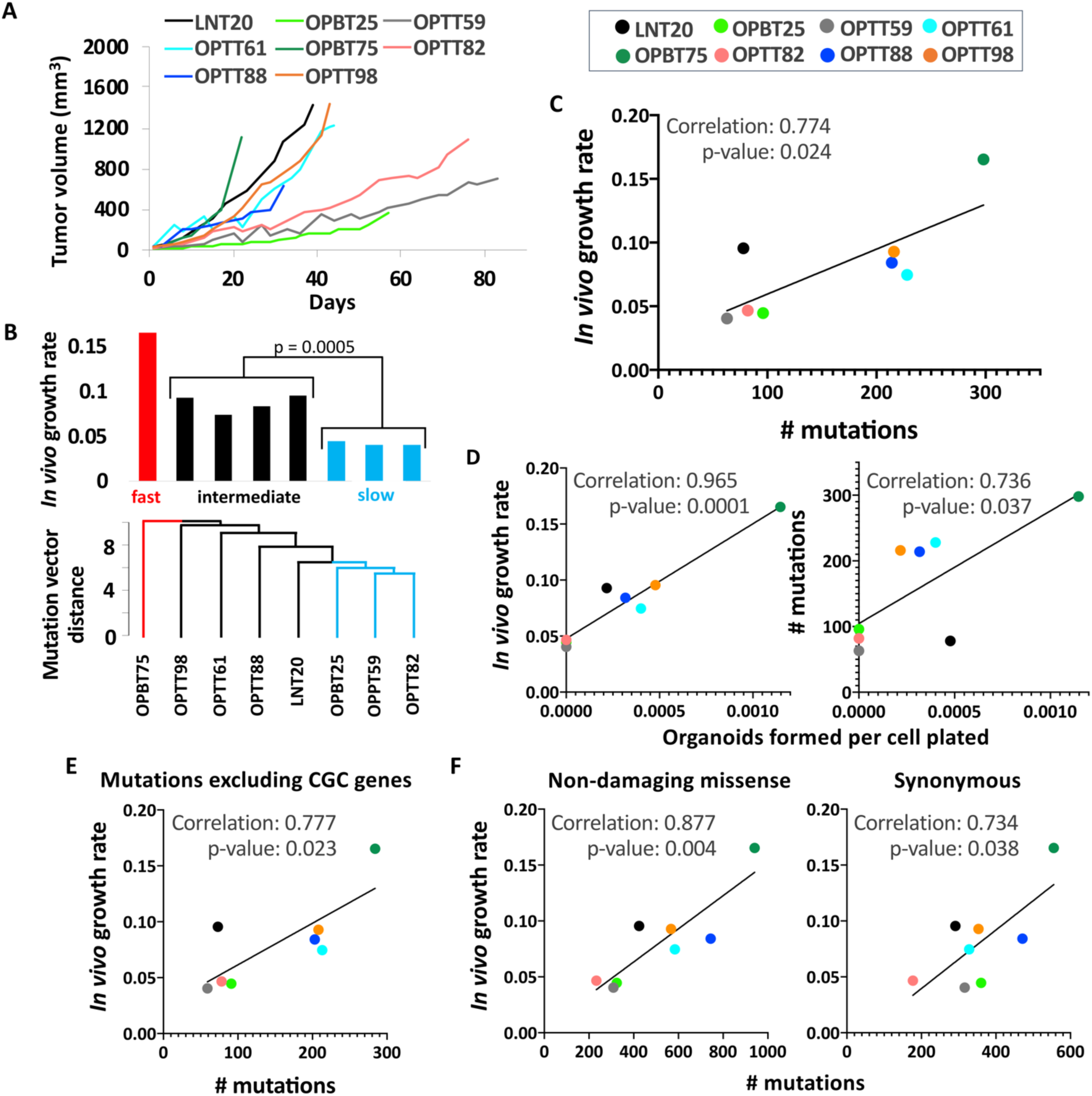
High mutational burden is associated with rapid growth of HPV+ PDXs and organoids. **A**. Representative *in vivo* growth curves of HPV+ PDXs during the first passage after initial engraftment **B**. Dendogram illustrating Euclidian distances among allele frequency-weighted mutation vectors in HPV+ PDXs (bottom). Growth rates of HPV+ PDXs are ordered by mutation vector distances (top). p-value was determined by Student’s t-test assuming unequal variances. **C**. *In vivo* PDX growth rates vs. total mutations per PDX. **D**. Organoid formation efficiency by PDX cells vs. *in vivo* growth rate (left) and number of mutations (right). **E**. PDX growth rates vs. number of mutations excluding all genes in CGC. **F**. PDX growth rates vs. substitutions excluded by mutation calling pipeline: missense mutations without damaging R-SVM scores (left) and synonymous substitutions (right). Pearson correlation coefficients and p-values based on two-tailed Student t-distribution are shown for each plot.

### High TMB is associated with HPV+ and HPV-tumor progression and is prognostic in early HPV-cases

The positive association between TMB and HPV+ PDX growth efficiency prompted assessment of TMB’s utility for predicting clinical behavior. Comparison of TMBs among 8th edition TNM stage groups for HPV+ HNSCCs identified decreased TMB in cases with limited (T1) local disease (Figure 4A, left). This finding was validated using the cervical squamous cell carcinomas in TCGA as a second HPV+ cohort (Figure 4A, right). To evaluate prognostic relevance of TMB, its distribution among the HPV+ HNSCCs was scanned by ROC analysis, which identified a cut point of 103 that best segregates outcomes based on Youden’s Index (Supplemental Table S7); however, this TMB cut-point was not predictive of overall survival (OS) (Figure 4B). Because local disease burden is more strongly predictive of poor outcome in HPV-cases, the same analysis was performed on the HPV-HNSCC cohort in TCGA. Here, TMB was significantly elevated in both locally advanced (T3/T4) and advanced stage (III/IV) disease (Figure 4C). Moreover, the optimal TMB cut-point for this cohort (99.5, Supplemental Table S8) predicted decreased disease-specific survival (DSS) and OS among early stage but not advanced stage cases with high TMB (Figure 4D). This finding was validated using TCGA data for esophageal squamous cell carcinoma (ESCC), a second HPV-malignancy with comparable risk factors, tissue origin, histology, and genetic landscape (35). As observed in HNSCCs, a TMB cut-point in ESCC was found that predicted OS for early but not advanced cases (Supplemental Figure S5). Together, these results support an association between high TMB and local progression of all HNSCCs and establish a prognostic relationship between high TMB and decreased survival of early stage HPV-cases.

**Figure 4.**
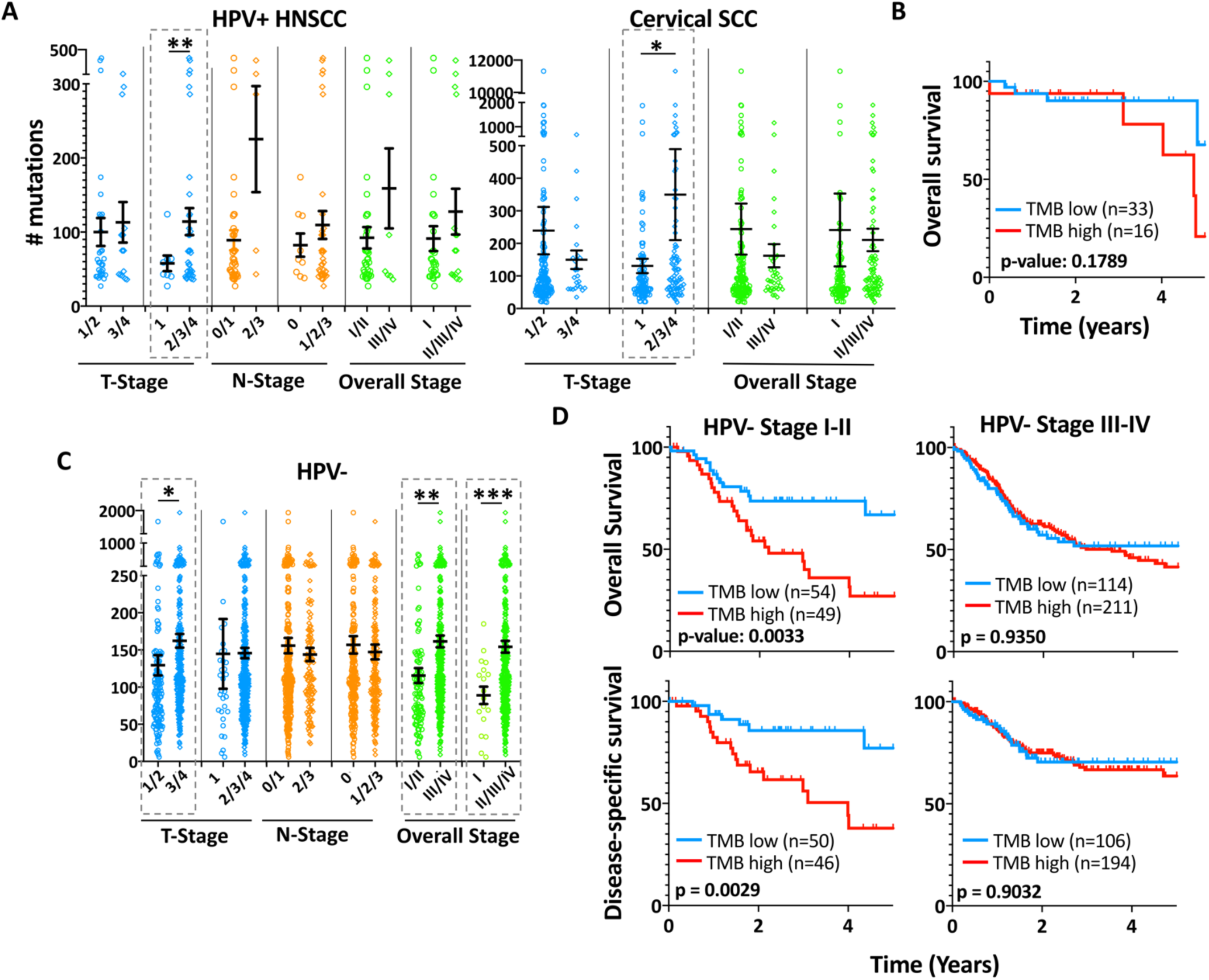
High TMB is associated with local progression in HPV+ and HPV-HNSCCs. Differences in mean TMB among different TNM and overall stage groups of HPV+ HNSCC (left) and cervical SCC (right) TCGA cases. *p<0.05, **p<0.025 Error bars represent SEM. p-values were determined by two-tailed Student’s t-test assuming unequal variances. **B**. OS for HPV+ HNSCCs in TCGA divided by a TMB cutoff of 103. **C**. Differences in mean TMB among different TNM and overall stage groups of HPV-TCGA cases. *p<0.05, **p<0.001, ***p<0.0001 **D**. 5-year OS and DSS for early stage (I/II) and advanced stage (III/IV) HPV-TCGA HNSCCs segregated by a TMB of 99.5. Staging for HPV+ HNSCC cases is updated to the 8^th^ edition AJCC clinical stage. 6^th^/7^th^ edition clinical stage is shown for HPV-cases. Survival p-values were determined by log-rank test.

### Reduced p16^INK4A^ levels in an atypical HPV+ PDX are shared by recurrent HPV+ cases in TCGA

Failure of TMB to aid risk stratification of HPV+ HNSCCs shifted focus to the molecular traits of the only HPV+ PDX whose patient of origin suffered an early recurrence, which had a lethal outcome (Supplemental Table S1). Among the tumor suppressors altered in this PDX, hemizygous loss of *CDKN2A* was a strikingly atypical feature relative to the other HPV+ PDXs and the 53 HPV+ HNSCCs in TCGA (Figure 5A). Presence of the hemizygous *CDKN2A* loss was further supported by DNA qPCR in the PDX and its tumor of origin (Figure 5B). Although the PDX and original tumor retained p16^INK4A^ overexpression by IHC, its p16^INK4A^ mRNA (Supplemental Figure S6) and protein levels (Figure 5C) were the lowest among the HPV+ PDXs. Because viral E7 acts upstream of p16^INK4A^ upregulation in HPV-related cancers, lower E7 was considered as a possible additional source of reduced p16^INK4A^ in this model. Among the PDXs, LNT20 also contained the lowest E7 transcript levels (Figure 5D), which was consistent with its relatively high levels of Rb protein (Figure 5E). Furthermore, LNT20 contained the highest ratio of phospho-Rb to Rb (Figure 5F), in keeping with the increase in CDK4/6 activity predicted by its lower p16^INK4A^ levels. To evaluate other aggressive HPV+ HNSCCs for related features, the HPV+ TCGA cases that recurred within 2 years of diagnosis were examined for p16^INK4A^ and E7 expression. These cases expressed significantly less p16^INK4A^ than the non-recurrent cases with equivalent follow-up (Figure 5G), despite complete absence of *CDKN2A* alterations in the HPV+ TCGA cohort. Although E7 transcript levels lacked correlation with p16^INK4A^ across the full range of p16^INK4A^ expression in the cohort (Figure 5H, left), a significant association was apparent at lower levels of p16^INK4A^ (Figure 5H, right). The findings were extended using RNA sequencing data for a second cohort of HPV+ oropharyngeal HNSCCs (36) (JHU), where p16^INK4A^ levels were reduced in the subset of cases that recurred (Figure 5I). Taken together, these findings identify an association between HPV+ HNSCC recurrence and reduced p16^INK4A^ upregulation downstream of lower viral E7 levels.

**Figure 5.**
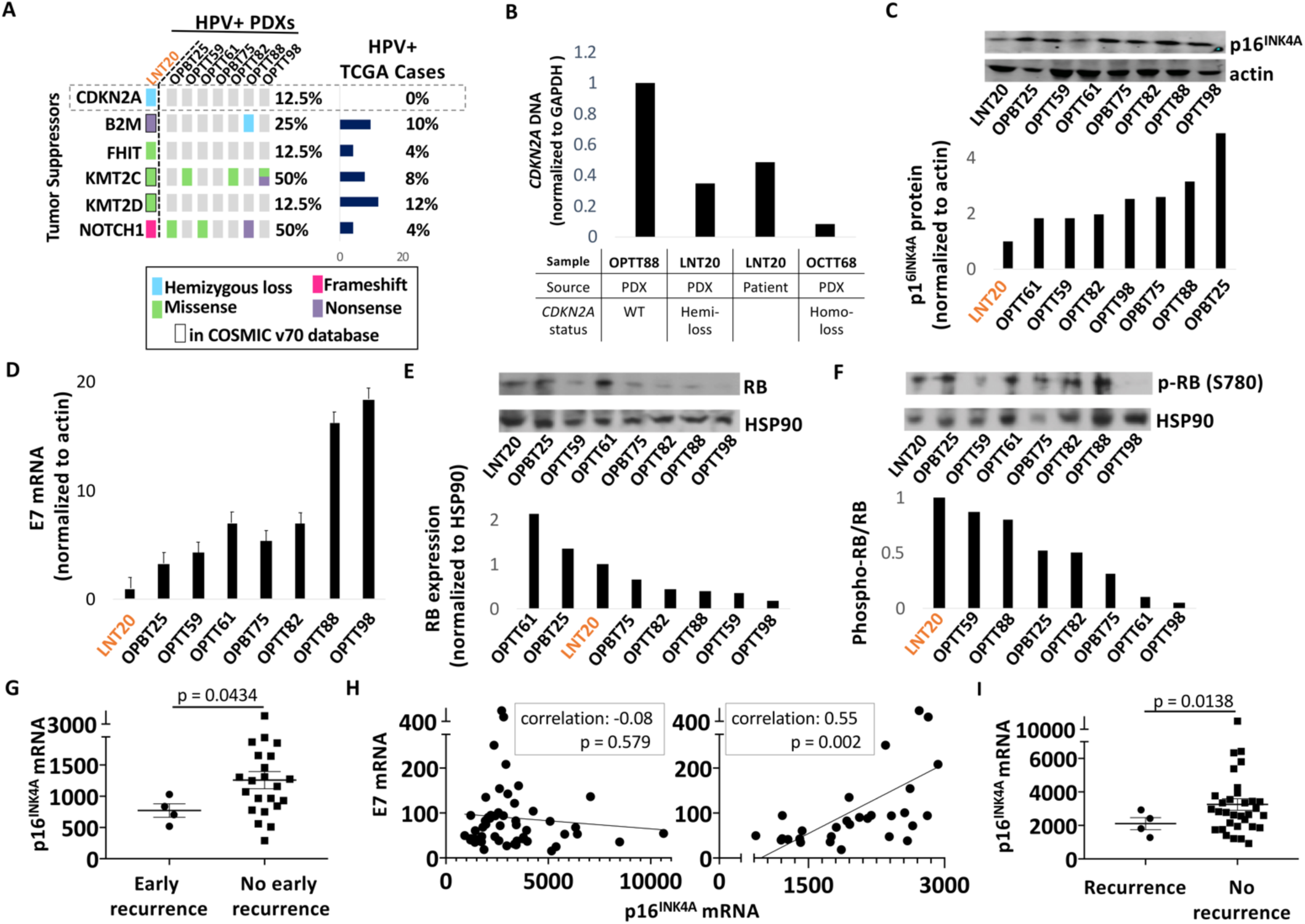
Molecular traits of an aggressive HPV+ PDX shared by recurrent HPV+ HNSCCs in TCGA. **A**. Tumor suppressors mutated or deleted in LNT20 relative to the other HPV+ PDXs and TCGA cases. **B**. *CDKN2A* DNA levels in the LNT20 PDX and its tumor of origin, relative to control PDXs with WT or deleted *CDKN2A*. **C**. p16^INK4A^ protein by western blot (top) with band densities normalized to actin (bottom) in HPV+ PDXs. **D**. HPV16 E7 mRNA levels normalized to actin in HPV+ PDXs. **E** RB protein (top) with corresponding band densities normalized to HSP90 (bottom) in HPV+ PDXs. **F**. phospho-RB protein (Ser780) (top) and phospho-RB to total RB ratios (bottom). **G**. p16^INK4A^ mRNA expression in TCGA HPV+ HNSCC recurred within 2 years (early recurrence) or were disease-free at the time of last early recurrence. **H**. E7 vs. p16^INK4A^ mRNA in all TCGA HPV+ HNSCCs (left) and those with a normalized p16^INK4A^ mRNA level of <3000 (right). Pearson correlation coefficients and p-values based on two-tailed Student t-distribution are shown for each plot. **I**. p16^INK4A^ expression in recurrent vs. non-recurrent HPV+ cases with matched follow-up duration in the JHU cohort. Bars represent mean ± SEM. p-values were determined by two-tailed Student’s t-test assuming unequal variances.

### The E2F target gene expression pattern is prognostic in HPV+ HNSCCs in two cohorts

Because E7 and p16^INK4A^ levels alone were not prognostic in TCGA, risk stratification was instead pursued by profiling expression of E2F target genes, which are upregulated via E7 in HPV+ HNSCCs and have lower expression in HPV-tumors (37). Hierarchical clustering of the 53 HPV+ oropharyngeal HNSCCs in TCGA using a list of 325 E2F target gene transcripts (37) identified a 22-case cluster (TCGA_C2) with decreased DFS and OS (Supplemental Figure S7). The target gene list was narrowed by excluding transcripts with area under the ROC curve (AUC) of <0.8 (Supplemental Table S9) to define 43 explanatory transcripts underlying TCGA_C2 (Figure 6A). Effective prediction of DFS and OS using the 43-transcript profile contrasted with the lack of prognostic utility for 8th edition AJCC staging in the same cohort (Figure 6B). Upregulation of these E2F target genes was diminished in TCGA_C2, where more genes were expressed at levels at least one absolute deviation below their medians for the total HPV+ case cohort (Figure 6C). Furthermore, most of the 43 E2F targets that were significantly upregulated in HPV+ vs. HPV-oropharyngeal HNSCCs in TCGA lacked upregulation in TCGA_C2 (Supplemental Figure S8). The case composition of TCGA_C2 was then compared to another molecular subgroup of HPV+ oropharyngeal cases in TCGA reported recently to have worse prognosis (7). In that study, 19 cases with worse survival outcomes were clustered based on attenuation of a 38-gene expression profile that best distinguishes HPV+ from HPV-HNSCCs in TCGA. Despite minimal overlap of those 38 genes with E2F target genes, the case cluster was found to be nearly identical to TCGA_C2 (Figure 6D), indicating diminished E2F target expression to be part of a wider decrease in viral effects upon host gene expression in poor prognosis cases. Validation of these results was pursued in another cohort containing 47 HPV+ HNSCCs (JHU) (30,36), which were clustered using the 43 prognostic E2F target transcripts identified from TCGA. In the JHU cohort, markedly worse DFS and OS were evident in a group of cases captured by hierarchical clustering (JHU_C2, Figure 6E), despite less variability in E2F target transcript levels across this cohort. Similar to TCGA_C2, JHU_C2 contained fewer transcripts with elevated expression relative to medians for the entire cohort (Supplemental Figure S9). Detailed clinical annotation in the JHU cohort further allowed determination that the E2F transcript profile was prognostic independent of whether patients received surgical or nonsurgical treatment (Supplemental Figure S10). Together, these results provide evidence that HPV+ HNSCCs with poor prognosis have reduced E2F target gene upregulation, which may be part of a broader decrease in the impact of the virus upon host gene expression.

**Figure 6.**
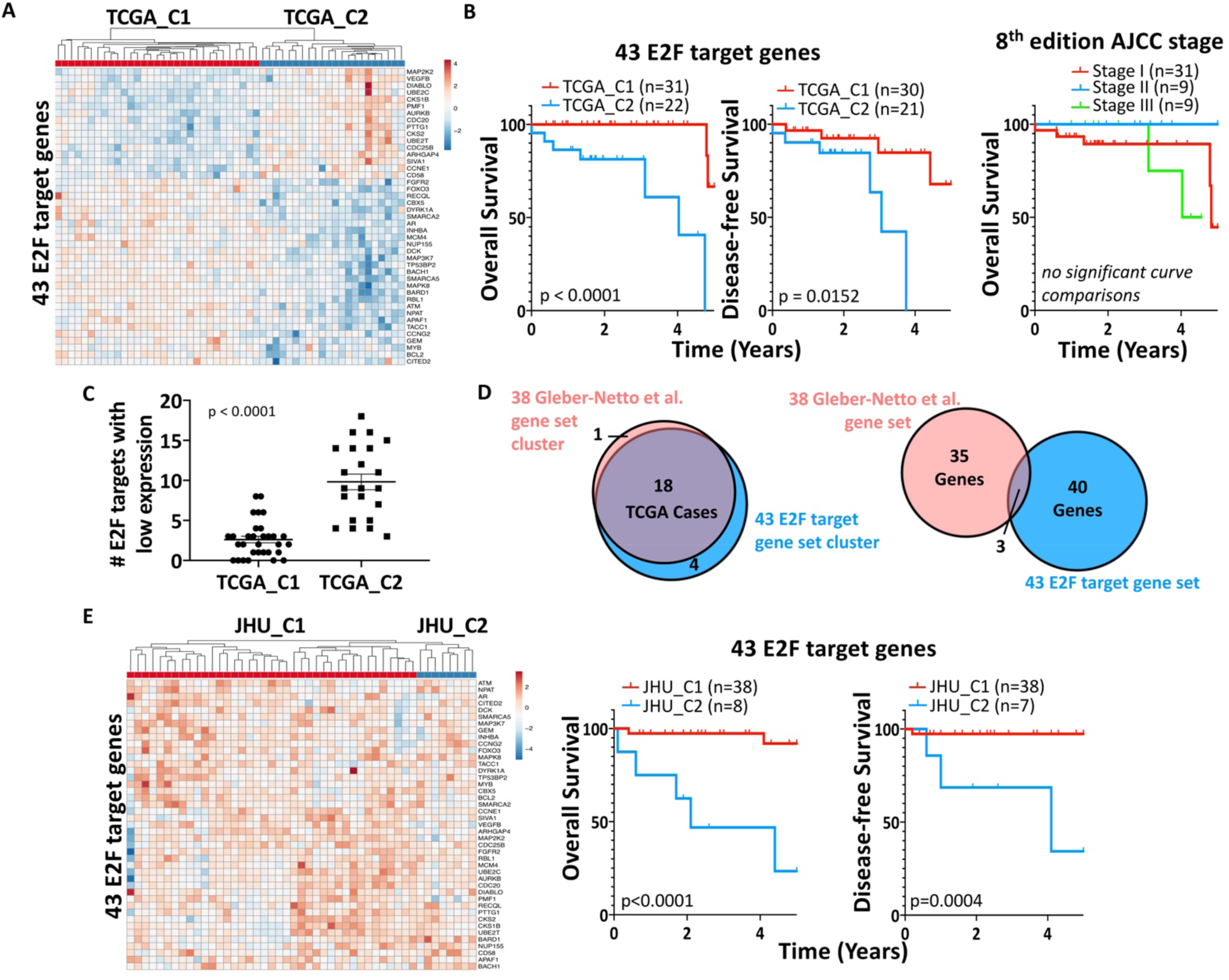
The E2F target gene expression pattern is prognostic in HPV+ HNSCCs in two cohorts. **A**. Expression of the 43 E2F target genes with AUC ≥0.8 in the HPV+ HNSCCs in TCGA. **B**. 5-year OS (left) and DFS (middle) for E2F target-derived clusters TCGA_C1 vs. TCGA_C2. OS of HPV+ TCGA HNSCCs segregated by 8^th^ edition clinical AJCC stage (right). **C**. Number of E2F target genes with low expression in TCGA_C1 vs. TCGA_C2 cases. p-value determined by two-tailed Student’s t-test assuming unequal variances. **D**. Overlap between HPV+ TCGA cases in TCGA_C2 and negatively prognostic cluster defined using 38 genes by Gleber-Netto et al. (left). Overlap of 43 E2F targets with the Gleber-Netto et al. gene set (right). **E**. Expression of 43 predictive E2F target genes among HPV+ HNSCCs in the JHU cohort (left). 5-year OS (middle) and DFS (right) for clusters JHU_C1 vs JHU_C2. Survival p-values were determined by log-rank test.

## DISCUSSION

This study addresses a need for improved preclinical resources to advance therapy for HPV+ HNSCC and simultaneously establishes a new basis for prognostic and therapeutic biomarker development. The HPV+ PDXs characterized here retained major genetic hallmarks of HPV+ HNSCC, including those lost in HPV+ cell lines (11). Despite potential for low PDX engraftment rates to create selection biases, the only bias identified was an over-representation of *NOTCH1* mutations and other Notch pathway alterations with predicted loss of function. An apparent enrichment of *NOTCH1* mutations by engraftment here is consistent with evidence for more aggressive clinical behavior of *NOTCH1*-mutant HPV+ HNSCCs (10). The models thus offer unique tools to address the biologic and therapeutic significance of attenuated Notch signaling in a tumor subtype that may have high clinical importance. A novel association in this study between local progression of the HPV+ HNSCC models and high TMB was preserved across multiple HPV+ and HPV-squamous cell carcinoma patient cohorts, including cases in the head and neck, esophagus, and cervix. However, TMB’s prognostic utility was limited to early stage HPV-tumors, and outcomes prediction for HPV+ HNSCCs was better served by the distinctive features of an HPV+ PDX from a rapidly lethal case. The reduced p16^INK4A^ upregulation downstream of low E7 levels in this outlier PDX was also observed in the HPV+ HNSCCs that recurred in two published cohorts. The decrease in E2F target gene expression predicted to accompany low E7 was shown to be prognostic in the same two cohorts, providing a new biologic basis to distinguish subgroups of HPV+ HNSCC patients warranting distinct therapeutic approaches.

The negative prognostic significance of reduced E2F target upregulation in this study adds to emerging evidence for oncogenic HPVs creating a greater imprint upon the host transcriptional landscape in the HPV+ HNSCCs that are readily curable. For instance, the poor prognosis HPV+ cases identified by us in TCGA using E2F target transcripts were nearly identical to a case cluster independently defined based on reduced HPV-driven dysregulation of a distinct set of genes (Figure 6D) (7). Such evidence that multiple molecular traits are widely shared among atypical HPV+ cases that recur after definitive therapy holds promise for future development of clinically useful risk stratifiers based on levels of a limited set of host and/or viral proteins.

Although the reduced p16^INK4A^ and E7 levels that prompted analysis of E2F target transcripts here lacked independent prognostic utility, they are intimately related to E2F target gene expression and might contribute to the prognostic transcriptional profile in some cases. Specifically, the low HPV E7 levels seen in the PDX and some early recurrent TCGA cases may reflect greater reliance on host oncogenic drivers and serve as one of multiple mechanisms of limiting E2F target upregulation in high-risk tumors. Similarly, the p16^INK4A^ upregulation thought to occur downstream of E7 expression in HPV-related cancers (38) was reduced in only a subset of recurrent cases in addition to the atypical PDX. In such instances, secondary events that prevent a reported addiction of HPV+ cancers to high p16^INK4A^ levels (39) may be necessary for cancer progression.

Because high p16^INK4A^ has potential to impair DNA repair by inhibiting homologous recombination (40), reduced p16^INK4A^ levels might also directly mediate the lethal phenotype of some HPV+ HNSCCs. A relative reduction in p16^INK4A^ levels in oropharyngeal HNSCCs without loss of clinical p16+ status may contribute directly to therapy failures by increasing resistance to radiation and cisplatin. This mechanism also makes the hemizygous *CDKN2A* deletion in the atypical HPV+ PDX and its lethal tumor of origin particularly intriguing. Although *CDKN2A* mutation or loss is commonplace and negatively prognostic in HPV-HNSCCs (41), these alterations are absent in HPV+ TCGA cases. However, *CDKN2A* alterations were recently reported in 10% of aggressive pulmonary metastatic recurrences from HPV+ HNSCCs (42), leading speculation that occasional *CDKN2A* alterations in untreated cases have negative prognostic significance and indicate poor candidates for therapy de-escalation.

Whereas the association between TMB and PDX model growth here did not have prognostic relevance in HPV+ HNSCCs, the reduced survival of early HPV-cancers with more mutations justifies further evaluation of TMB in that clinical context. High TMB was linked to local progression in both HNSCC subtypes, but lesser ability for HPV+ local disease to reduce survival might explain prognostic utility being limited to HPV-disease. To date, high TMB has been extensively studied as a predictor of immunotherapy responses mediated by neo-epitope abundance in a tumor (43-46) but not as a correlate of the tumor-autonomous progression quantified here in the models. In this regard, prediction of rapid tumor growth in the milieu of an immunodeficient mouse is a novel finding that may lead to expanded utility for TMB as a biomarker. Rapid progress of TMB assays toward clinical application (47) thus may create opportunities to simultaneously identify certain high-risk tumors based on TMB at presentation and escalate initial therapy for them via checkpoint inhibition.

Over-representation of *NOTCH1* mutations in the HPV+ models adds to existing evidence that PDX generation sometimes selects for aggressive tumor genotypes (15, 17-19). The loss-of-function mutants in the Notch pathway in HPV+ and HPV-HNSCCs indicate a tumor suppressor role (22) and have previously been associated with poor outcomes in HPV-cases (48). Canonical Notch signals in normal squamous epithelia promote hierarchical differentiation (49) and their loss enhanced tumorigenesis in a HNSCC mouse model expressing E6/E7 (33). Furthermore, Notch signal activation downstream of E2F function is observed in development (50) and served to limit tumor growth in a hepatocellular carcinoma model upon Rb loss (51). Thus, silencing Notch may similarly cooperate with E7-mediated Rb degradation to drive carcinogenesis in select HPV+ HNSCCs.

There are notable limitations to the insights gained here from the models and their utility as preclinical tools. First, the small size of the panel does not capture the full genetic diversity among HPV+ HNSCCs. Secondly, relatively high cure rates in this disease led to the PDXs capturing only one lethal case, thus limiting the models’ capacity to represent the most aggressive HPV+ tumors. Of note, all patients of origin for the PDXs completed aggressive multimodality therapy, and thus PDX engraftment may still have enriched for tumor phenotypes at risk of recurrence during ongoing therapy de-escalation efforts for this disease. Third, *NOTCH1* mutation has been shown to be prognostic in only one HPV+ HNSCC cohort thus far (10). Despite that study being the largest case series linking HPV+ HNSCC mutations to outcomes, lack of adequately powered validation cohorts presently makes the significance of the Notch pathway alterations in the PDXs less clear. Finally, the prognostic expression profile defined here using 43 E2F target genes could not be easily reduced to a single transcript that is prognostic across multiple cohorts, whereas doing so would greatly facilitate further development of a clinical biomarker. Nevertheless, the substantial knowledge gained from this small HPV+ PDX panel illustrates the value of pursuing patient-derived models even for tumor types that do not grow efficiently outside humans.

## MATERIALS AND METHODS

### Patient derived xenografts and organoids

PDXs were established, passaged, and cryopreserved as described by us (13) using NOD/SCID/IL-2Rγ^-/-^ mice under University of Pennsylvania IRB protocol #417200 and Wistar Institute IACUC protocol #112652. Tumor volumes were calculated from width^2^*length/2. PDX growth rates were estimated by the slope from a linear regression model for the logarithm of tumor volume as a function of time for each replicate. Growth rates were then calculated from the inverse-variance weighted sum of the replicates for each PDX. PDXs were dissociated as described (13) and resuspended in Matrigel (600 cells/μL) for organoid culture. 50μl was plated in 24-well format with 500μL of organoid media. Organoid media was comprised of DMEM/F12^+^ (Gibco, Gaithersburg, MD) supplemented with N2, B27, 50 ng/mL recombinant human EGF (Thermo Fisher Scientific, Waltham, MA), 0.1 mM *N*-acetyl-L-cysteine (Sigma, St. Louis, MO), 2% Noggin/R-Spondin–conditioned media, 50 μg/mL Gentamicin (Invitrogen, Carlsbad, CA), Glutamax, 10 mM HEPES (Gibco), and 10 μM Y27632 (SelleckChem, Houston, TX).

### Sequencing and Analysis

Whole exome sequencing of PDXs was performed from snap-frozen tissue using the Illumina HiSeq (2×150bp) platform. Genomic DNA extraction was performed with Qiagen QIAamp DNA Mini Kit (Qiagen, Hilden, Germany) per the manufacturer’s recommendation. DNA was quantified with Qubit 2.0 DNA HS Assay (Thermo) and quality-assessed by Tapestation genomic DNA assay (Agilent Technologies, Santa Clara, CA). Library preparation was performed using KAPA Hyper Prep kit (KAPA Biosystems, Wilmington, MA) per manufacturer’s instructions. Exome capture was performed with IDT xGen Exome Research Panel v1.0 (IDT, Skokie, IL). Library quality and quantity were assessed with Qubit 2.0 DNA HS Assay (Thermo), Tapestation High Sensitivity D1000 Assay (Agilent), and QuantStudio® 5 System (Applied Biosystems, Foster City, CA).

DNA for targeted sequencing was obtained from snap-frozen or formalin-fixed tissue, and *NOTCH1* and *PIK3CA* exons were sequenced using an Ion AmpliSeq Custom Panel (Thermo). DNA was isolated using the PureLink Genomic DNA Kit (Invitrogen) for snap frozen primary specimens and QIAamp DNA FFPE Tissue kit (Qiagen) for FFPE primary specimens. The Qubit Broad Range dsDNA kit (Thermo) and the NGS FFPE QC kit (Agilent, Santa Clara, CA) assays were used to determine DNA quantity and quality, respectively.

Deletions and damaging mutations (frameshift, stopgain, stoploss, missense with damaging R-SVM score (34)) were predicted to confer loss of function, while amplifications and non-damaging missense mutations in the COSMIC v70 database were considered gain-of-function. For principal component analysis (PCA), damaging mutations were weighted by allele frequency. Non-transformed continuous scores for each gene were used to calculate principal components.

### Mutational variant calling

Mutational variants were processed through the bcbio pipeline and mapped to the human reference genome (hg19) using Burrows-Wheeler Aligner (BWA). GATK HaplotypeCaller recalibrated the mapped reads based on quality and realigned them around indel regions. Low complexity regions (LCRs) were removed and mutational variants were called when identified by two out of four variant callers: Freebayes, Vardict, MuTect2, or VarScan. Variants were limited to exomic mutations with allele frequencies ≥4% and minimum read depths ≥10. In the absence of matched normal controls, somatic variants were identified if their population frequency was <1% based on the 1000 Genome Project, ESP6500, ExAC, and CG46 databases. Only nonsynonymous, frameshift, stoploss, or stopgain mutations were called. Nonsynonymous mutations were further filtered by either a damaging radial support vector machine (R-SVM) score or presence in the COSMIC v70 database. Finally, mutations that passed all other filtering criteria but were found in three or more PDXs were discarded due to potentially being missed by the population frequency analysis.

### Copy number analysis

Copy number alterations (CNAs) were called using the CNVkit software toolkit and mapped to the human reference genome (hg19) using Burrows-Wheeler Aligner (BWA). Each sample was first median-centered and corrected for read-depth bias. In the absence of matched normal controls, the threshold for calling amplifications or deletions was based on the scaled log2 copy ratios of each CNA segment relative to a generated ground reference. The ground reference was calculated by taking a weighted average (Tukey’s biweight location) of the log2 converted read-depth value of each bin across all samples. Thresholds for deletions and amplifications were log2 values of <-0.25 and >0.2, respectively.

### TCGA data analysis

The cBioPortal (52) was used to analyze genomic and clinical information from the Provisional head and neck(24) and esophageal (53), and PanCancer Atlas cervical (54) squamous cell carcinoma TCGA data sets. Additional patient data including clinical follow-up were downloaded from the Genomic Data Commons. The HPV+ oropharyngeal HNSCCs in TCGA were identified based on expression of viral E6/E7 (20). Overall and disease-free survival data were updated for TCGA HPV+ cases using follow-up tables V1.0 and V4.8, and cases with persistent disease were assigned disease-free survival of 1 day. HPV+ TCGA tumors were restaged clinically by the 8^th^ edition AJCC manual. Full clinical annotation of the 53 HPV+ oropharyngeal HNSCCs in TCGA is provided in Supplemental Table S2. Normalized mRNA expression for TCGA samples was downloaded from the Broad GDAC Firebrowse repository and ln(x+1)-transformed for clustering. Unsupervised hierarchical clustering and heatmap visualization were performed with ClustVis (55) using correlation distance and average linkage.

### Real-time quantitative PCR (qPCR)

*CDKN2A* copy number was estimated by qPCR as described (56). Briefly, PCR reactions were carried out in a volume of 20 µl, using 10 μl of 2X *Power*SYBR Green PCR Master Mix (Applied Biosystems). The thermal cycling conditions were: 95°C for 10 min followed by 40 cycles at 95°C for 15 s, 60°C for 30 s and 72°C for 30 s. The primers used were: CDKN2A forward: 5’-GGCTGGCTGGTCACCAGA-3’; CDKN2A reverse: 5’-CGCCCGCACCTCCTCTAC-3’; GAPDH promoter forward: 5’-TACTAGCGGTTTTACGGGCG-3’; GAPDH promoter reverse: 5’-TCGAACAGGAGCAGAGAGCGA-3’. Reverse transcriptase qPCR was performed as previously described (13). Primers for reverse transcriptase qPCR were: HPV16 E7 forward: 5’-CCGGACAGAGCCCATTACAA-3’; HPV16 E7 reverse: 5’-CGAATGTCTACGTGTGTGCTTTG-3; HPV16 E6 forward: 5’-TCAGGACCCACAGGAGCG-3’; HPV16 E6 reverse: 5’-CCTCACGTCGCAGTAACTGTTG-3; E2 forward: 5’-TGGAAACACATGCGCCTAGAA-3’ E2 reverse: 5’-GTTGCAGTTCAATTGCTTGTAATGC-3’; p16 forward: 5’-AGCATGGAGCCTTCGGCTGA-3’; p16 reverse: 5’-CCATCATCATGACCTGGATCG-3’.

### Western Blotting and Immunohistochemistry

PDX tissues were lysed in RIPA buffer (Thermo) and protein quantified by BCA assay (Thermo). 50ug protein was separated on 10% ECL gels (GE, Pittsburgh, PA) and transferred to nitrocellulose using the Trans-Blot® System (Bio-Rad, Hercules, CA). Antibodies are listed in Supplemental Table S10.

### Statistical analysis

Mutational frequencies among cohorts were compared by Fisher’s exact test. Differences in mutation frequency, TMB, and expression among cohorts were evaluated using Student’s t-test assuming unequal variances. Significance testing of Pearson correlations was performed using Student’s t-test. Survival was compared by log-rank test.

## AUTHOR CONTRIBUTIONS

NDF and DB developed the concept, designed experiments, and wrote the manuscript.

NDF acquired data and performed data analysis and interpretation.

PR, ATP, KTM, and JJ performed data analysis and interpretation.

VS, CDJ, BW, IMM acquired data and contributed to data interpretation.

PAG performed data analysis, contributed to study design, and provided critical review of the manuscript.

FOGN, JAC, CRP provided material support and critical review of the manuscript.

GSW, AL, AKR, HN, JAC, CRP, ELW, BW, IMM, and RBC provided conceptual advice and critical review of the manuscript.

## ACKNOWLEDGEMENTS

This work is supported by NIH R01-DE027185 (D Basu, P Gimotty), P30-DK050306 (Core Facilities), P30CA016620 (Bioinformatics Core of the Abramson Cancer Center), and P30CA010815 (Wistar Institute Cancer Center).

