## Supplemental Information for "Identifying predictors of HPV-related head and neck squamous cell carcinoma progression and survival through patient-derived models"

Supplemental Figures S1-S10

Supplemental Tables S1-S8, S10

**SUPPLEMENTAL FIGURES**

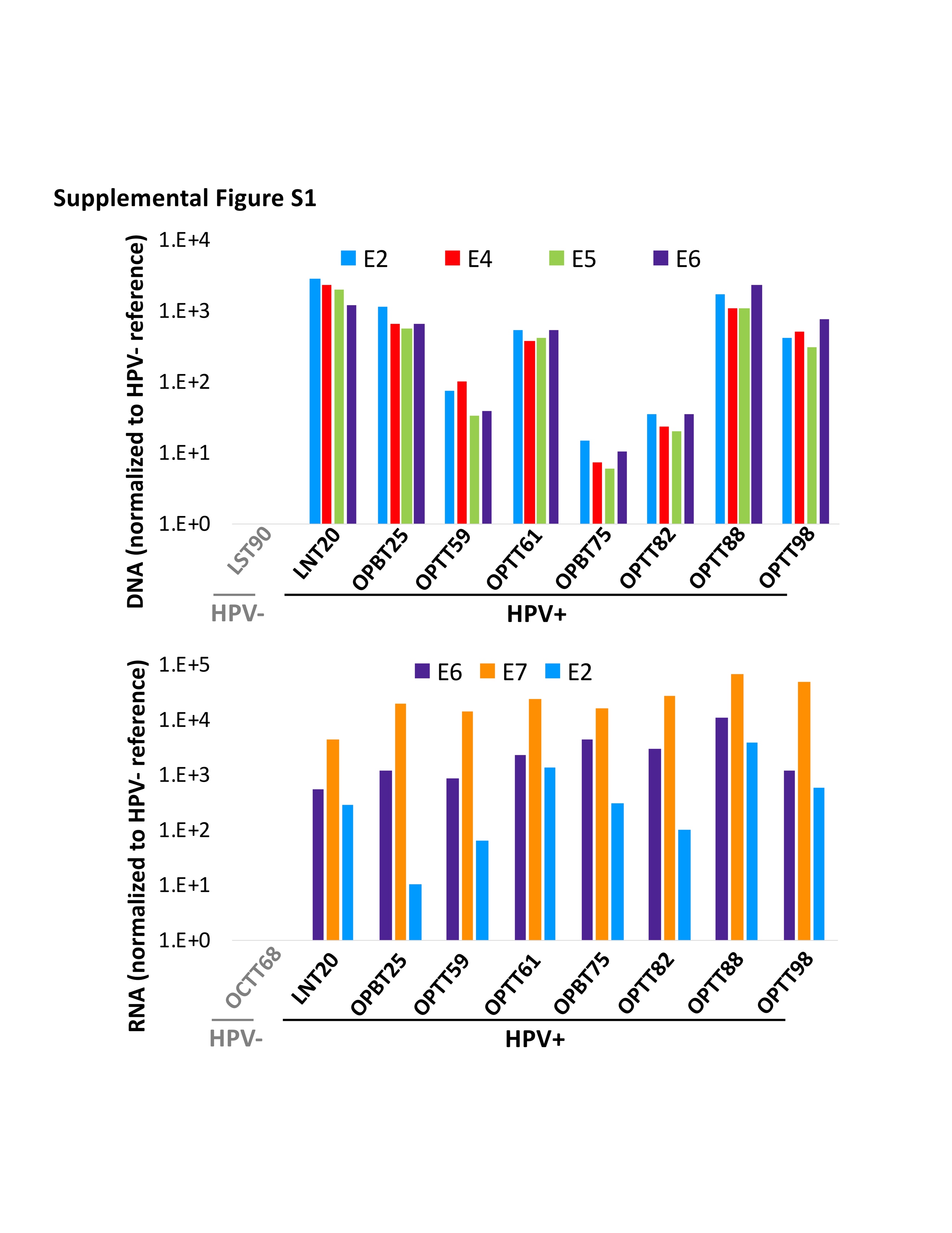

**Supplemental Figure S1. HPV16 viral DNA and RNA detected in HPV+ PDXs.** HPV16 viral oncogene DNA (top) and RNA (bottom) levels by qPCR in HPV+ PDXs normalized to GAPDH and actin, respectively, and represented relative to HPV- reference PDX.

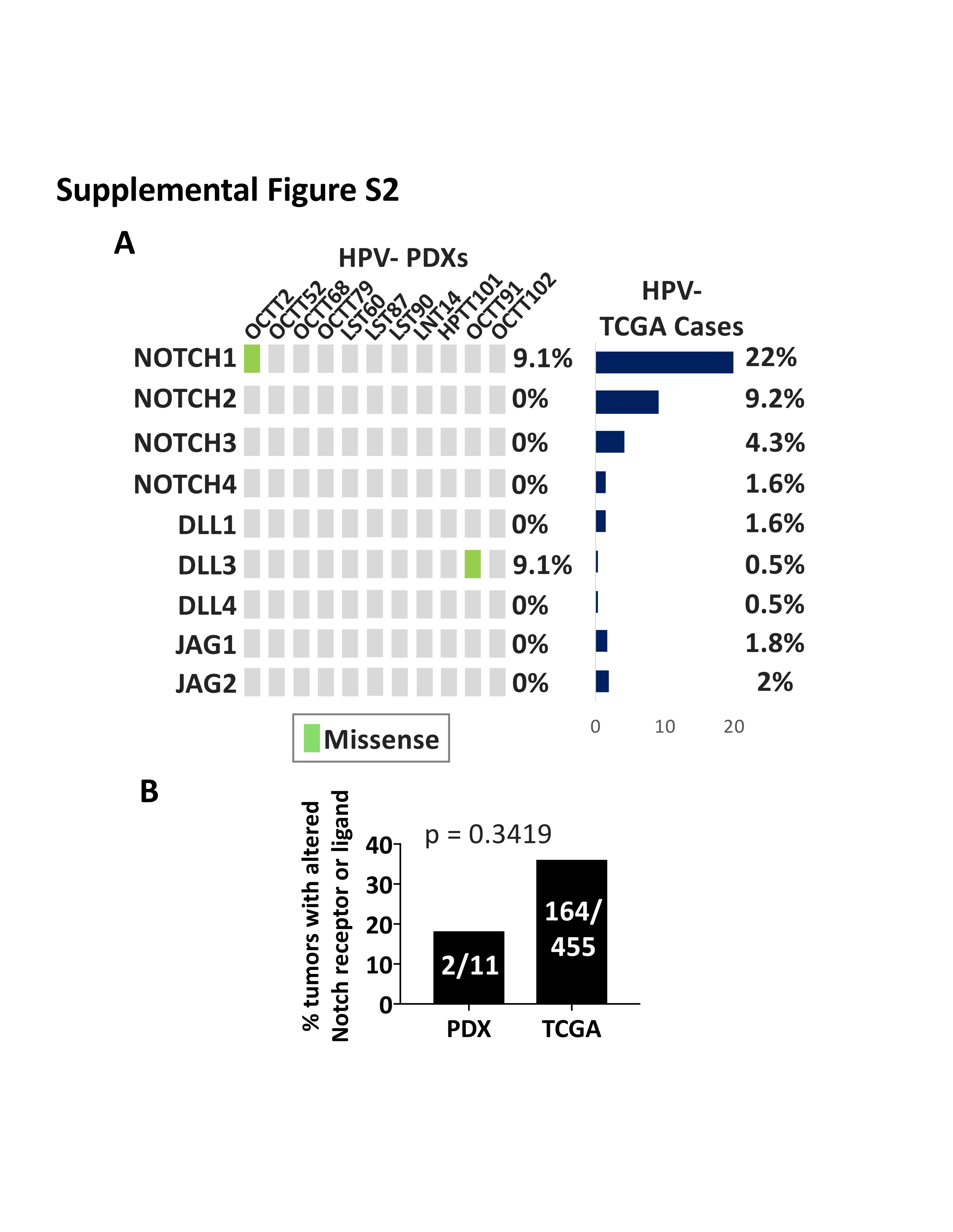

**Supplemental Figure S2. Notch pathway mutations and CNAs in HPV- PDXs and TCGA cases.** **A.** Predicted loss-of-function mutations and CNAs in Notch receptors and ligands in HPV- PDXs and TCGA cases. **B.** Frequency of mutations and/or CNAs in NOTCH receptor or NOTCH ligand genes in HPV- PDXs vs. HPV- TCGA cases. p-value determined by Fisher’s exact test.

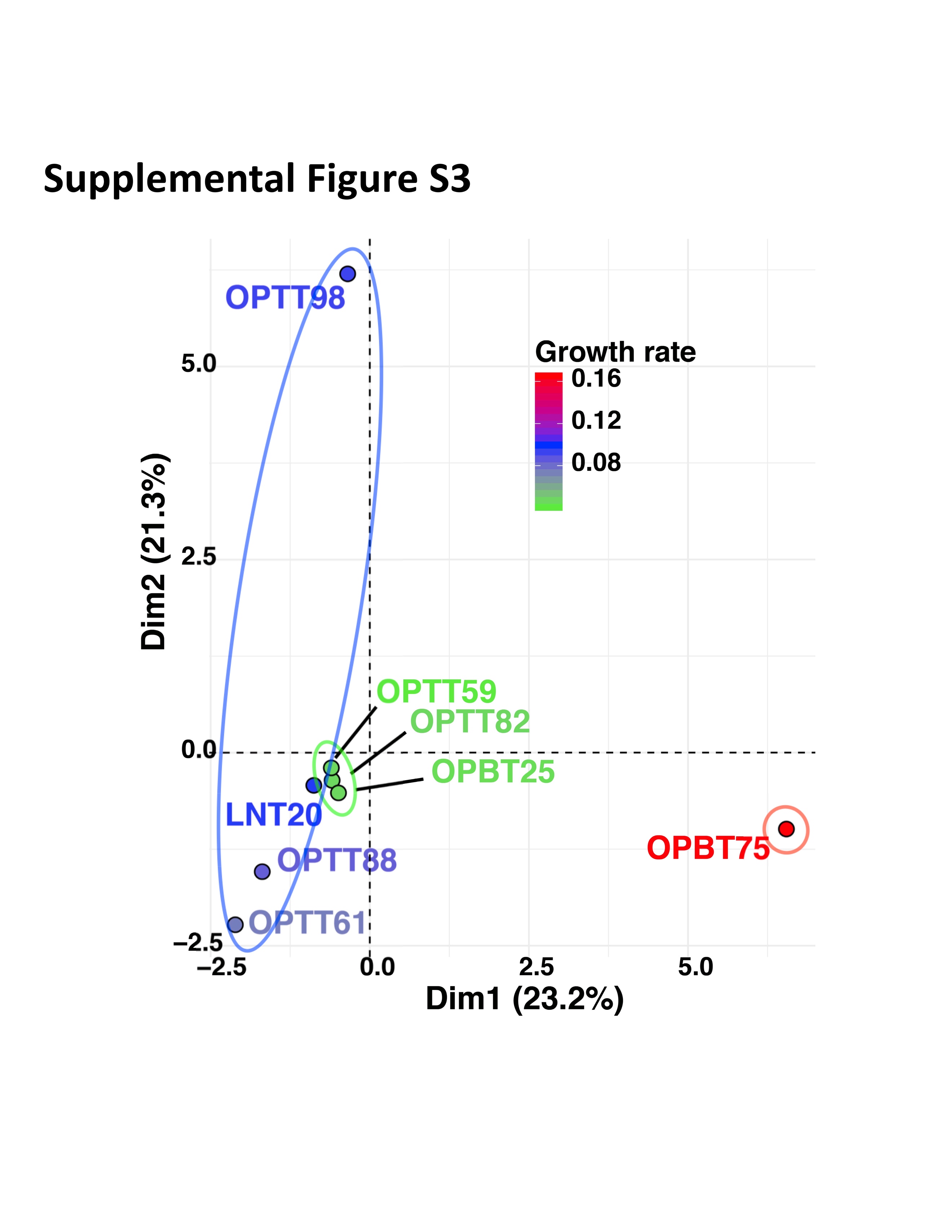

**Supplemental Figure S3. PCA projection of mutation profile vs. growth rate in HPV+ PDXs.**

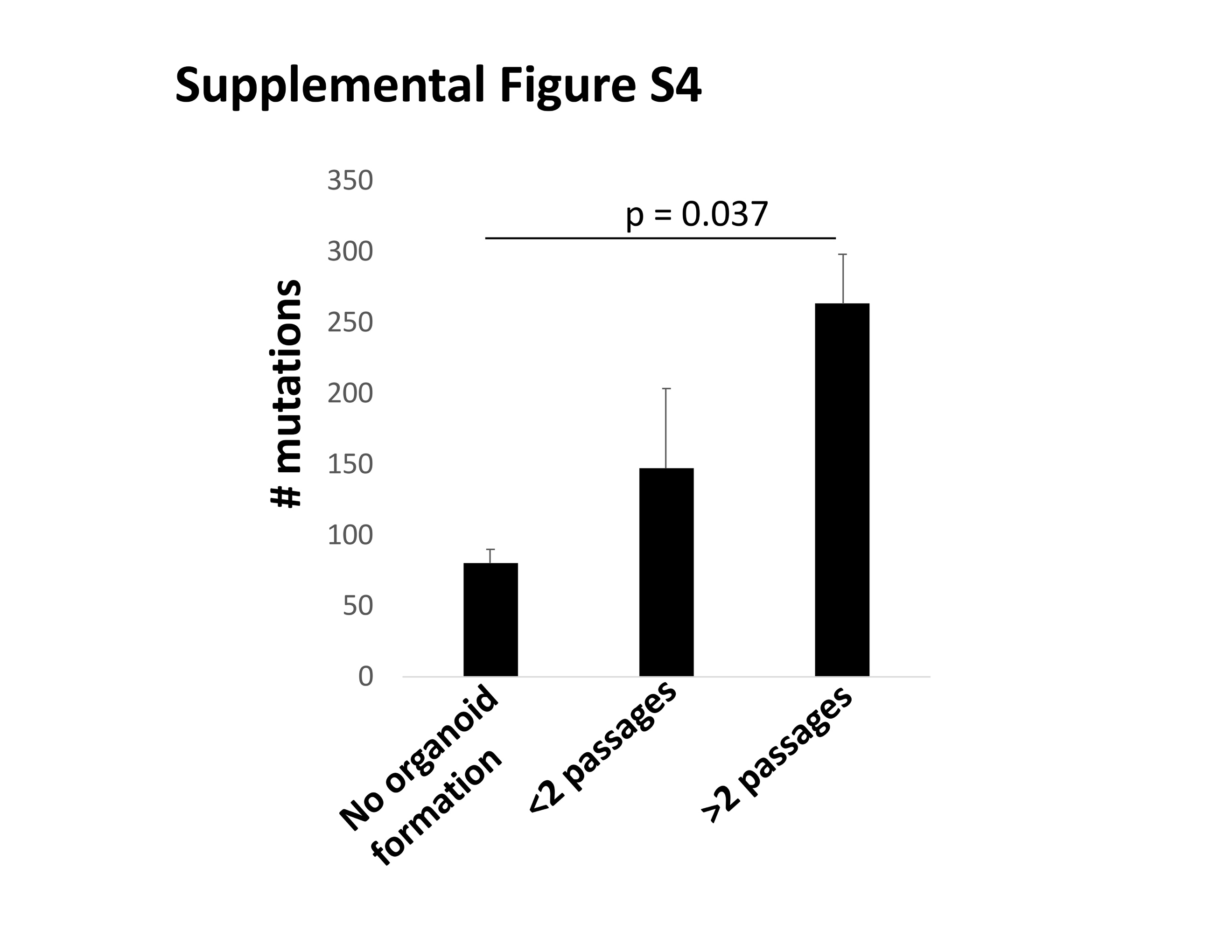

**Supplemental Figure S4. HPV+ PDXs with high *in vitro* organoid formation potential have high TMB.** TMB of HPV+ PDXs that do not form organoids, form organoids that cannot be stably passaged (<2 passages), and form organoids that can be continuously passaged *in vitro* (>2 passages). p-value determined by two-tailed Student’s t-test assuming unequal variances.

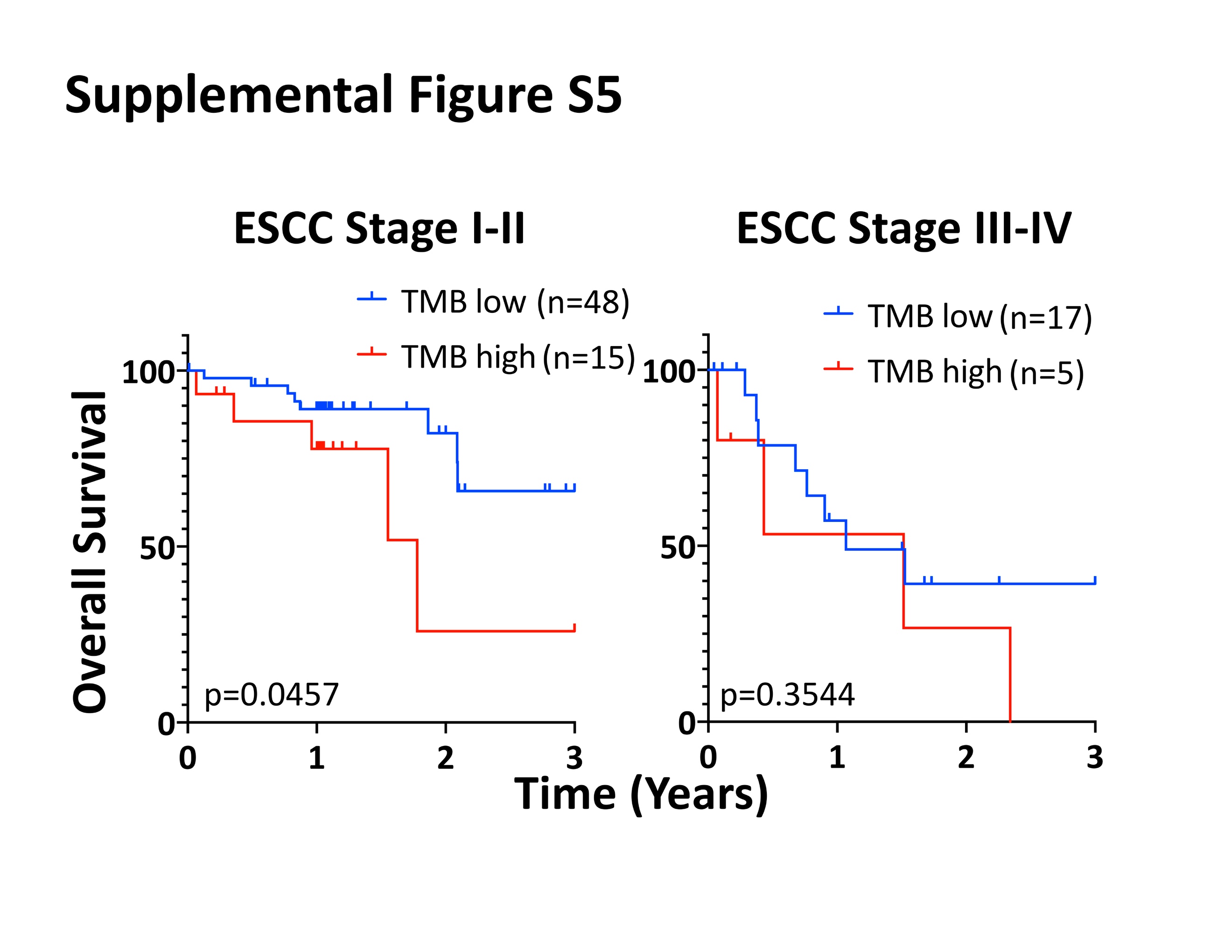

**Supplemental Figure S5. TMB is associated with survival in early stage ESCCs.** 3-year OS for early stage (I/II) and advanced stage (III/IV) TCGA ESCC cases segregated by a TMB of 198. Log-rank p-values are inset. Staging is 6^th^/7^th^ edition AJCC.

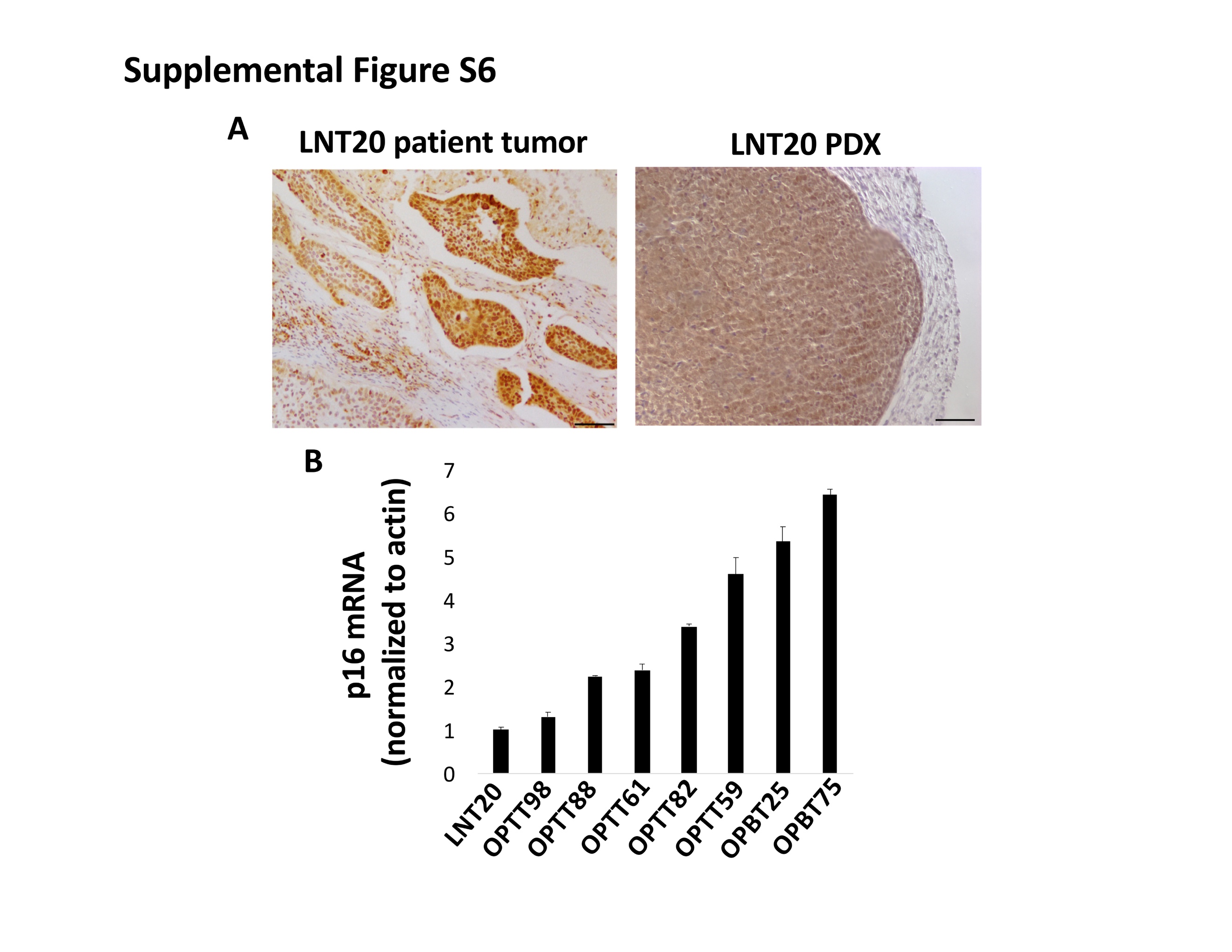

**Supplemental Figure S6. p16^INK4A^ overexpression is retained by LNT20 but is reduced relative to the HPV+ PDXs. A.** IHC staining detects p16^INK4A^ in LNT20 patient specimen and PDX (10x, Bar=100µm). **B.** p16^INK4A^ mRNA levels normalized to actin in HPV+ PDXs.

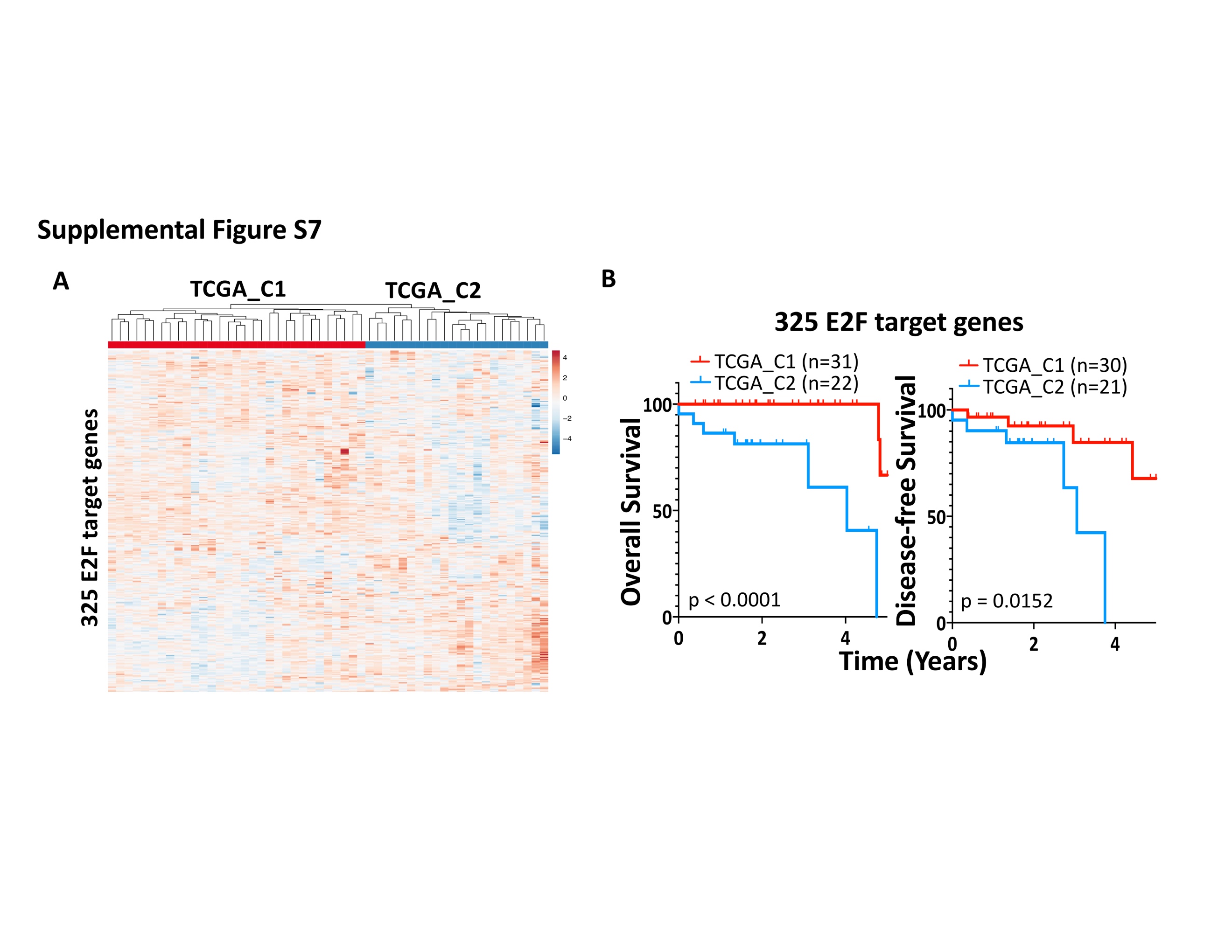

**Supplemental Figure S7. HPV+ TCGA cases form 2 distinct clusters defined by E2F target gene expression.** **A.** Expression of 325 E2F target genes across the HPV+ HNSCCs in TCGA. **B**. 5-year OS and DFS for E2F target-derived clusters TCGA_C1 vs. TCGA_C2.

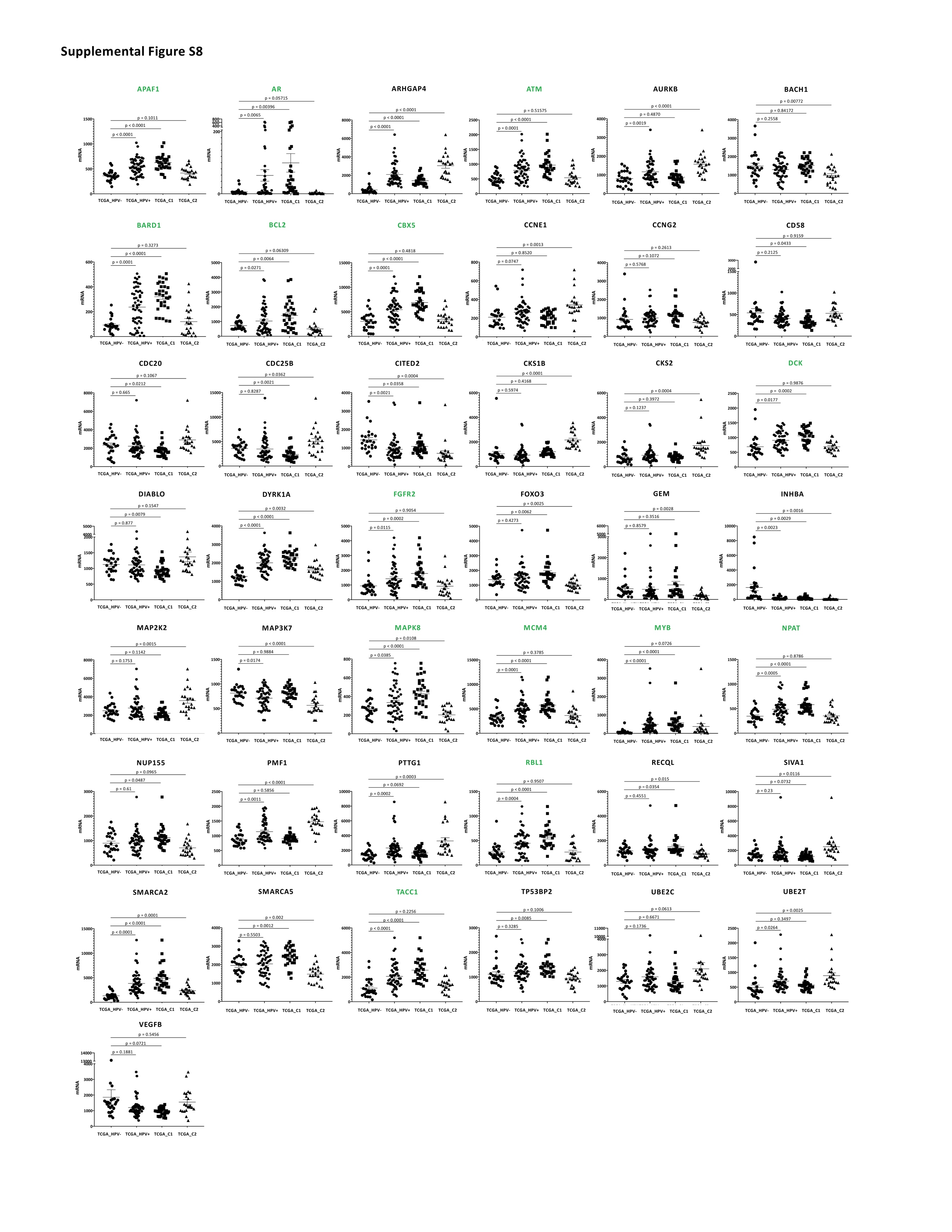

**Supplemental Figure S8. Majority of 43 E2F targets upregulated in HPV+ cases lose increased expression in TCGA_C2.** Comparison of mRNA levels of 43 E2F targets among TCGA cohorts: HPV- oropharyngeal (TCGA_HPV-), TCGA_HPV+, HPV+ cluster TCGA_C1, and HPV+ cluster TCGA_C2. Targets highlighted in green indicate genes with significantly increased expression in all 53 HPV+ cases compared with HPV- oropharyngeal cases but not in the subset of HPV+ TCGA_C2 cases. Bars represent mean ± SEM.

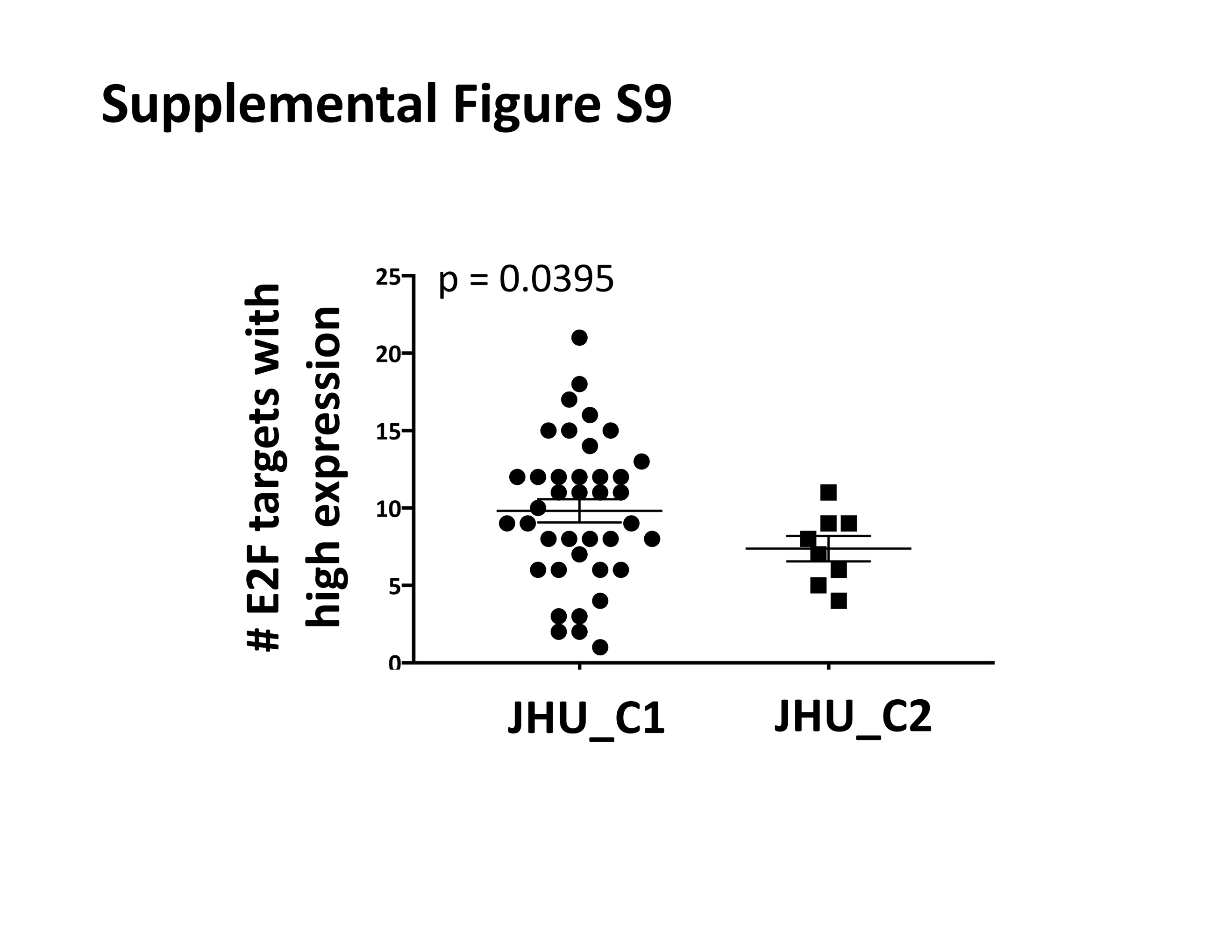

**Supplemental Figure S9. Cases in JHU_C2 have less E2F target upregulation.** Number of E2F target genes with high expression (more than 1 absolute deviation greater than median expression across all 47 JHU HPV+ HNSCCs) in JHU_C1 vs. JHU_C2 cases. p-value determined by two-tailed Student’s t-test assuming unequal variances.

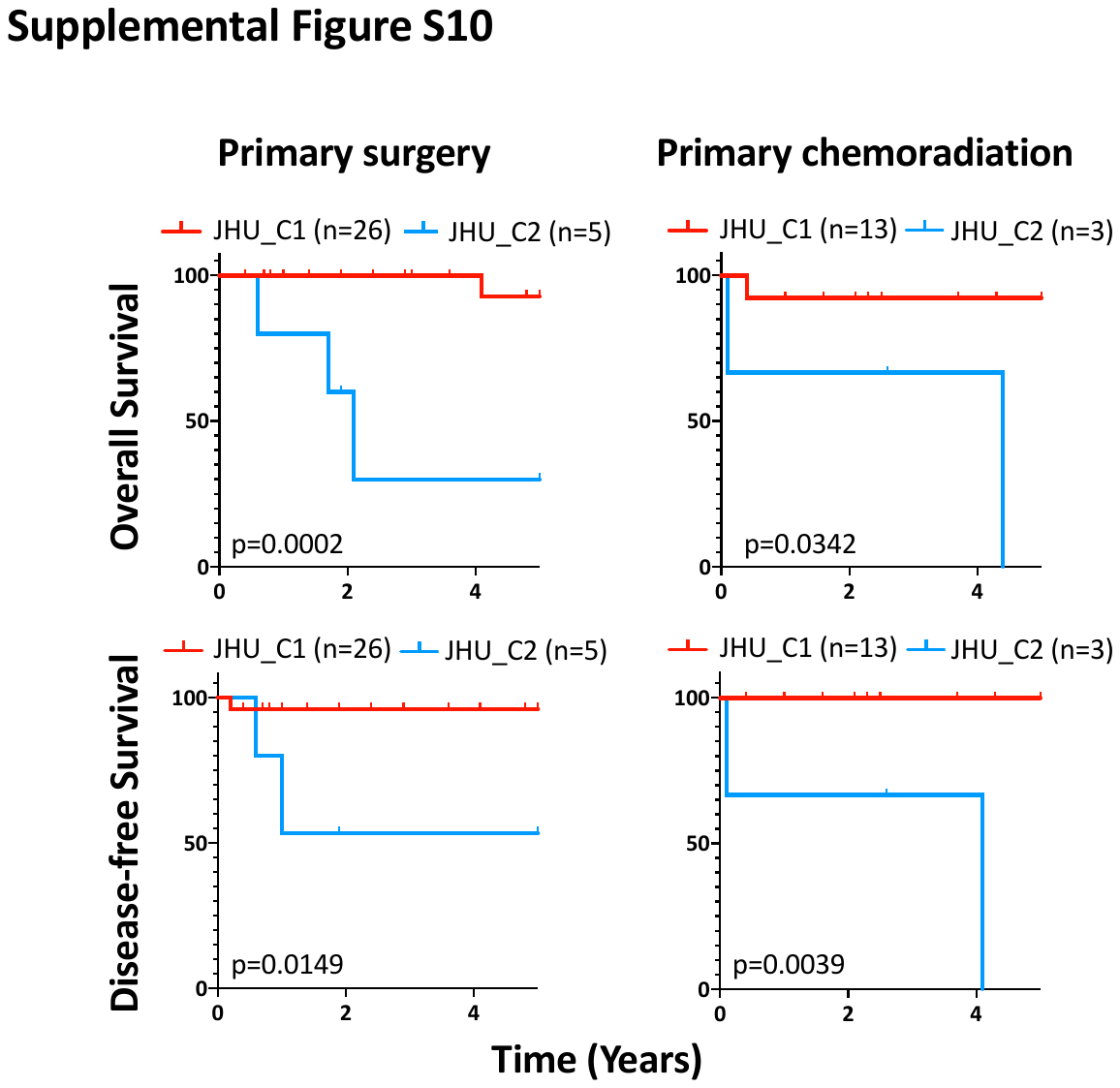

**Supplemental Figure S10. JHU_C2 cases that received either primary surgical or non-surgical definitive treatment have poor prognosis.** 5-year OS (top) and DFS (bottom) survival for clusters JHU_C1 vs JHU_C2 segregated by primary mode of treatment. Survival p-values were determined by log-rank test.

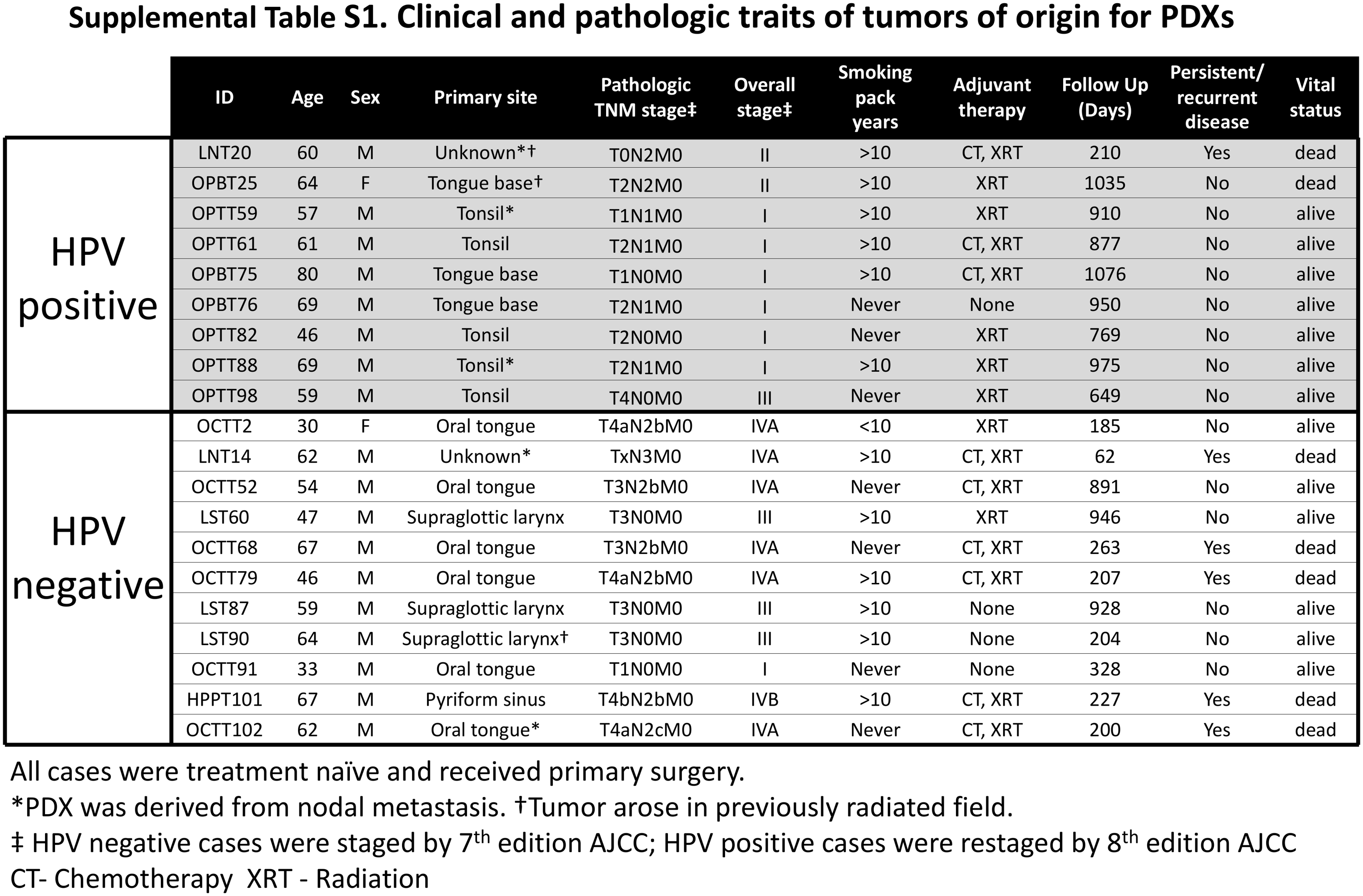
**SUPPLEMENTAL TABLES**

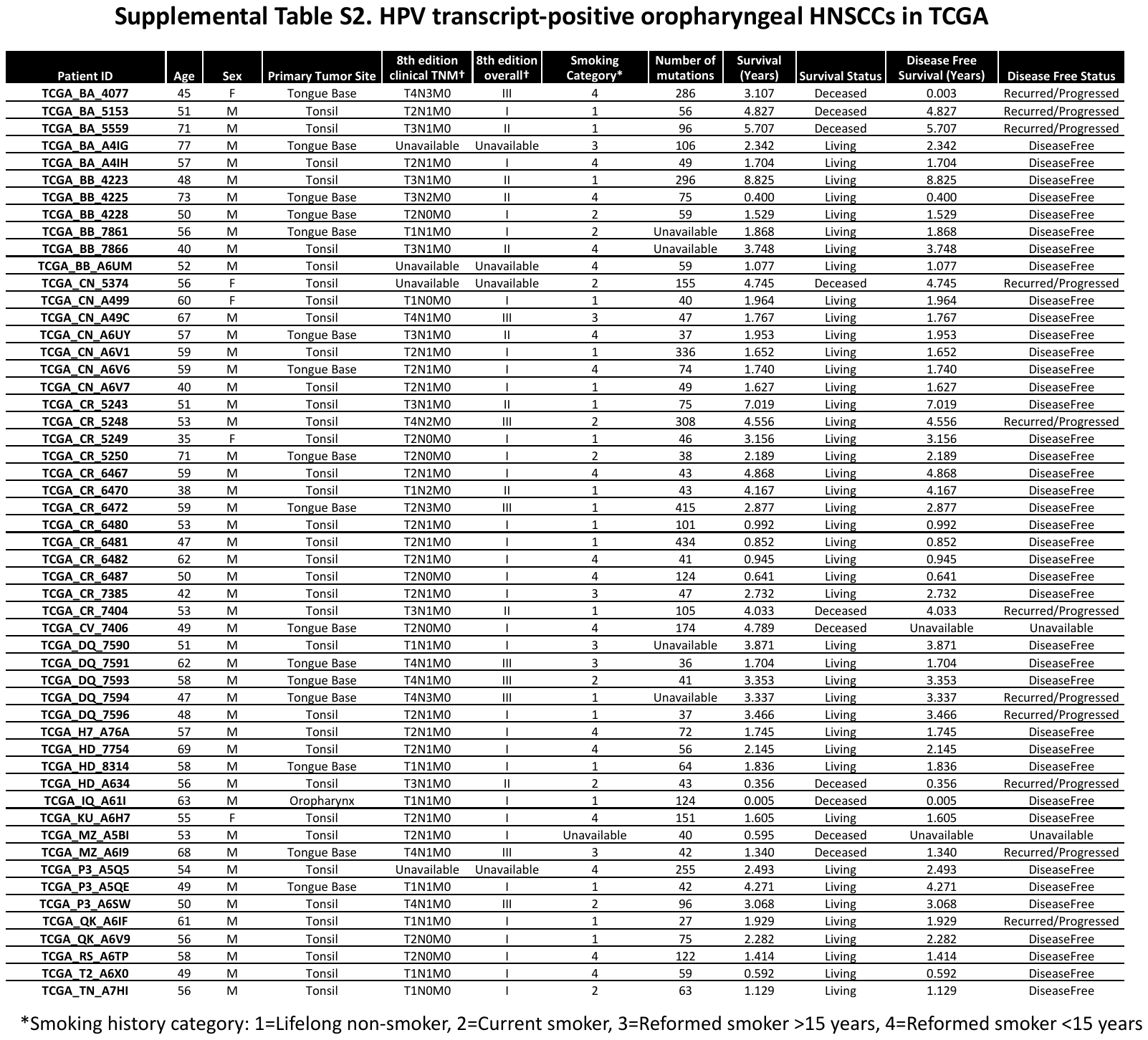

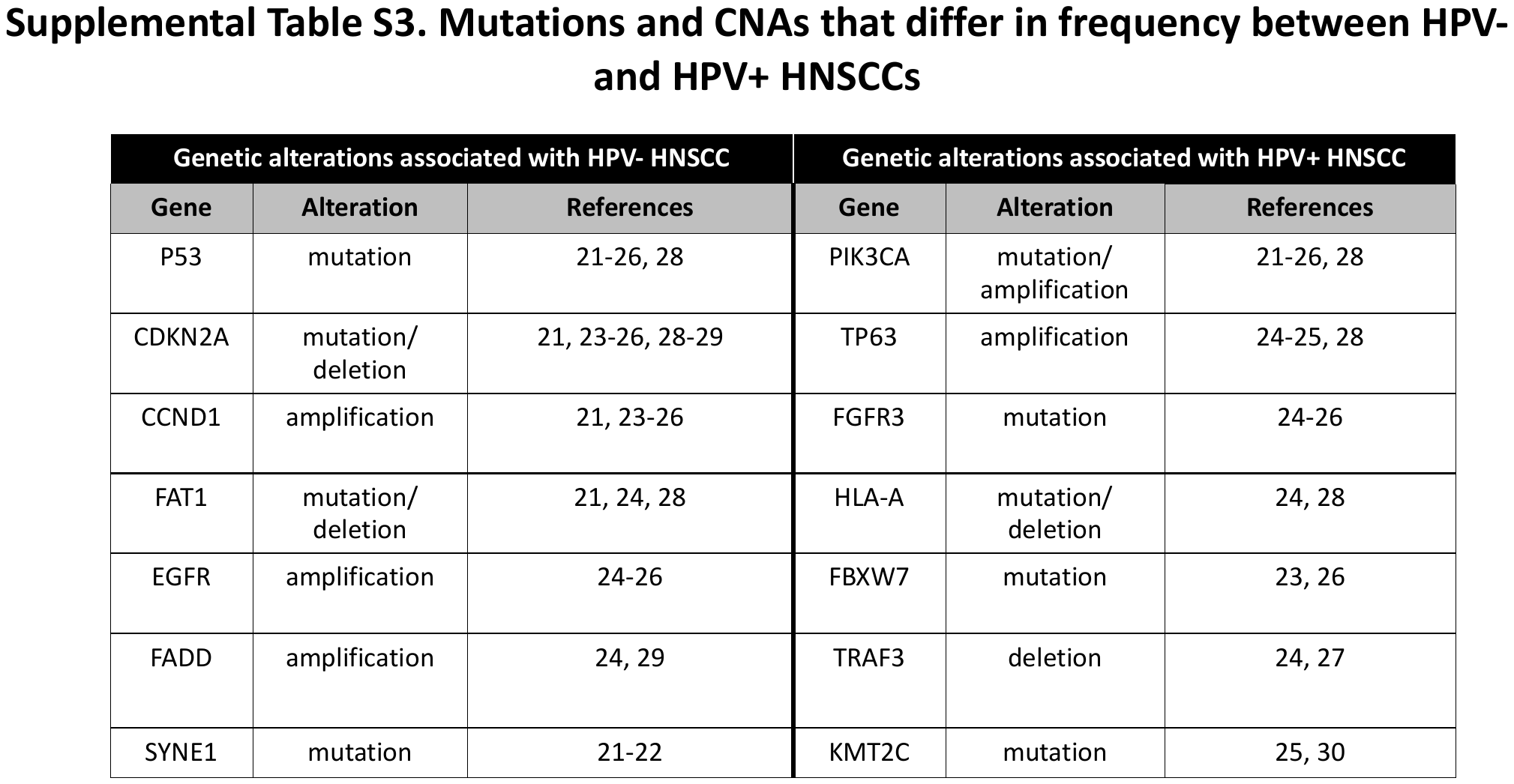

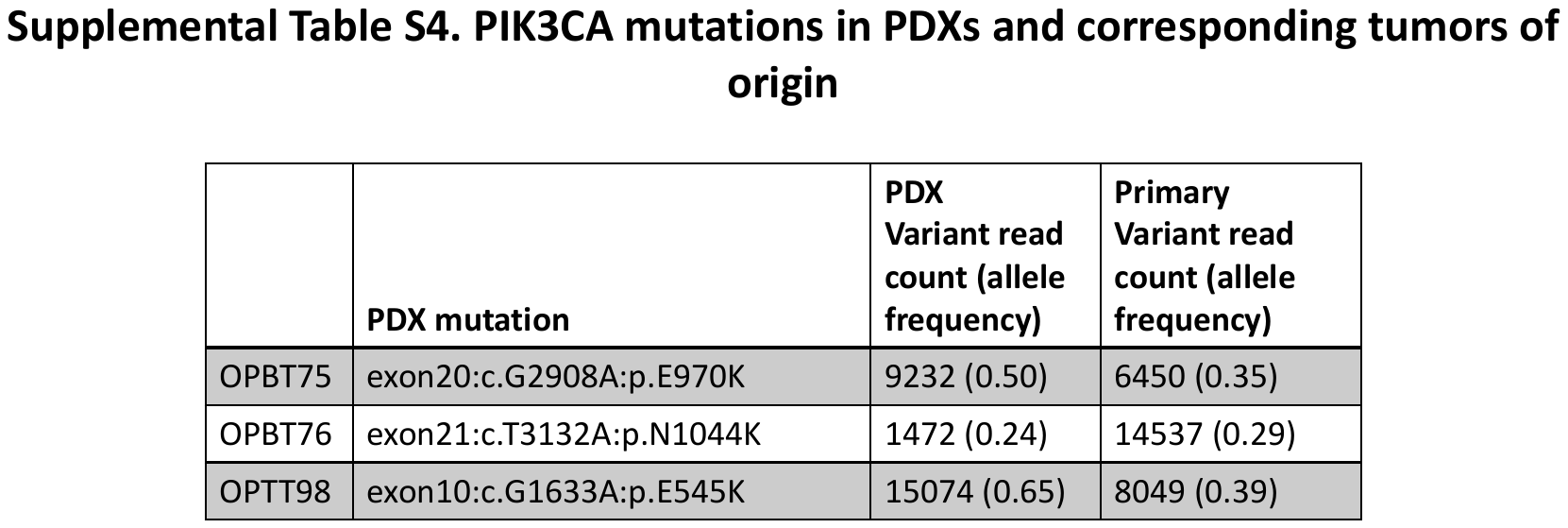

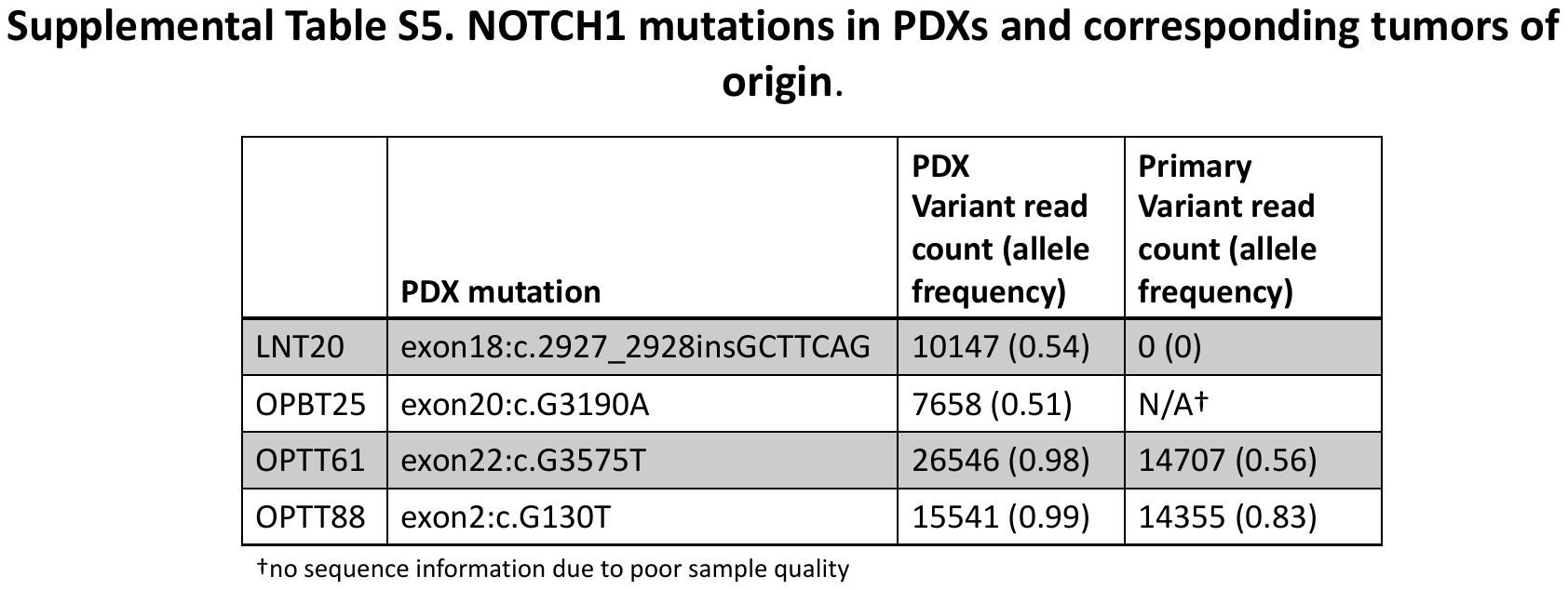

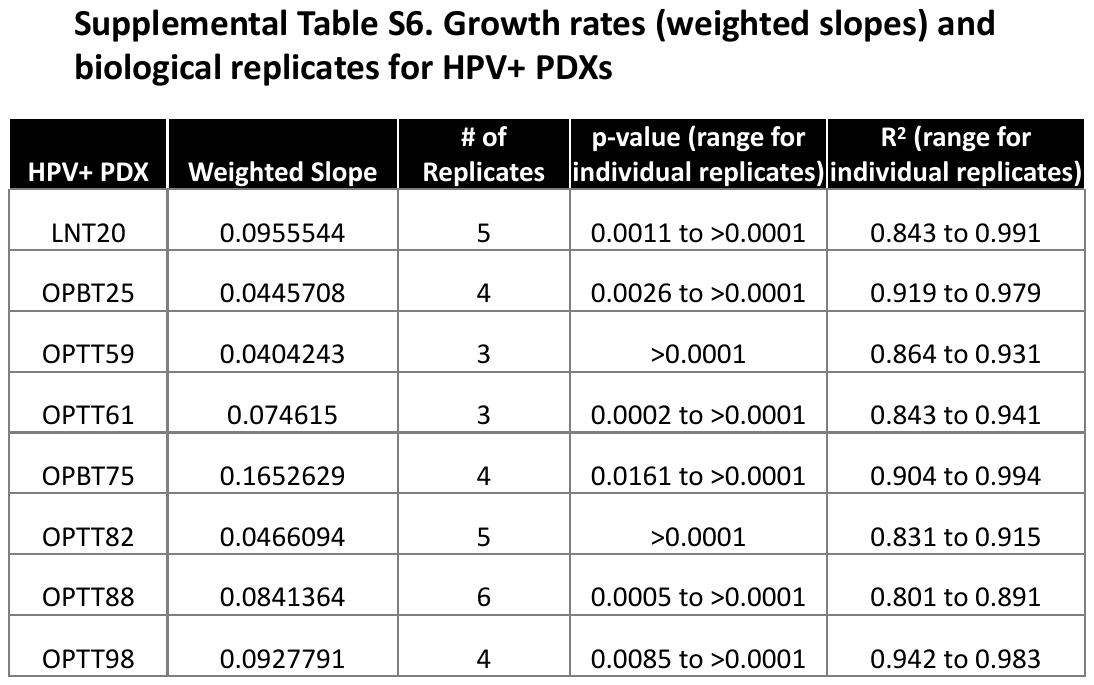

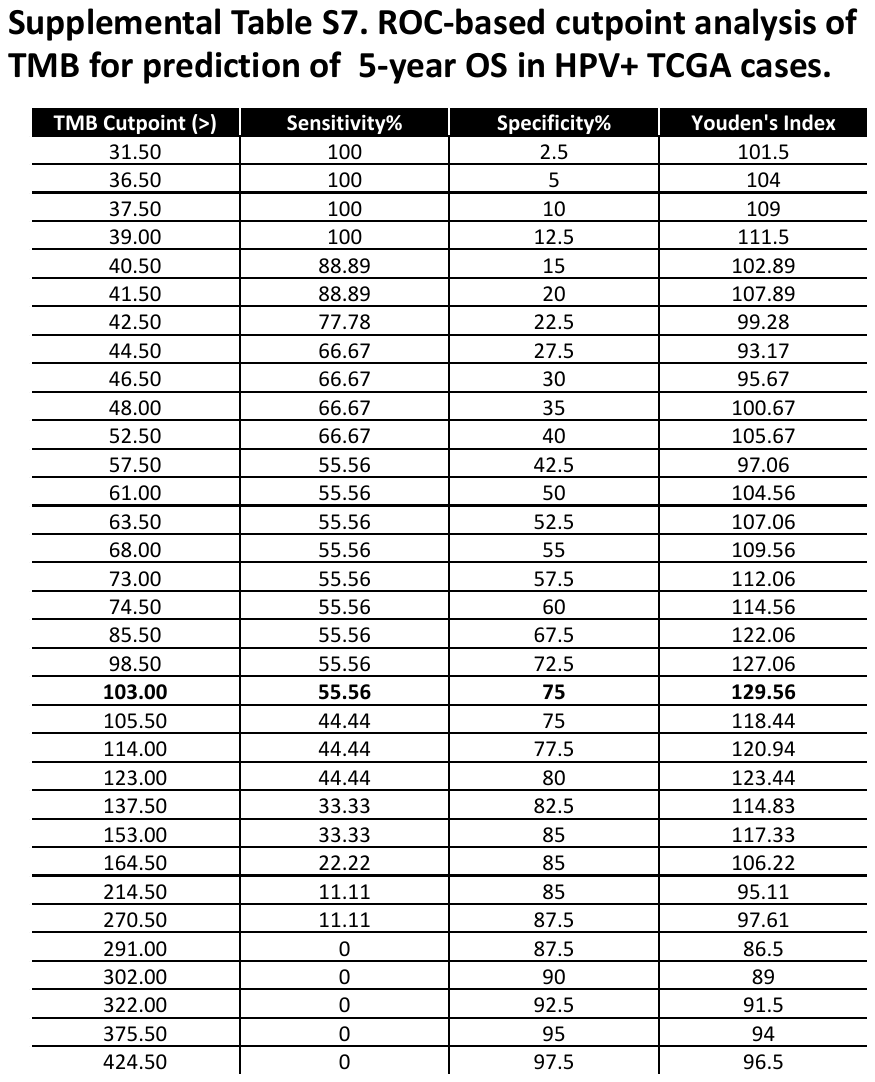

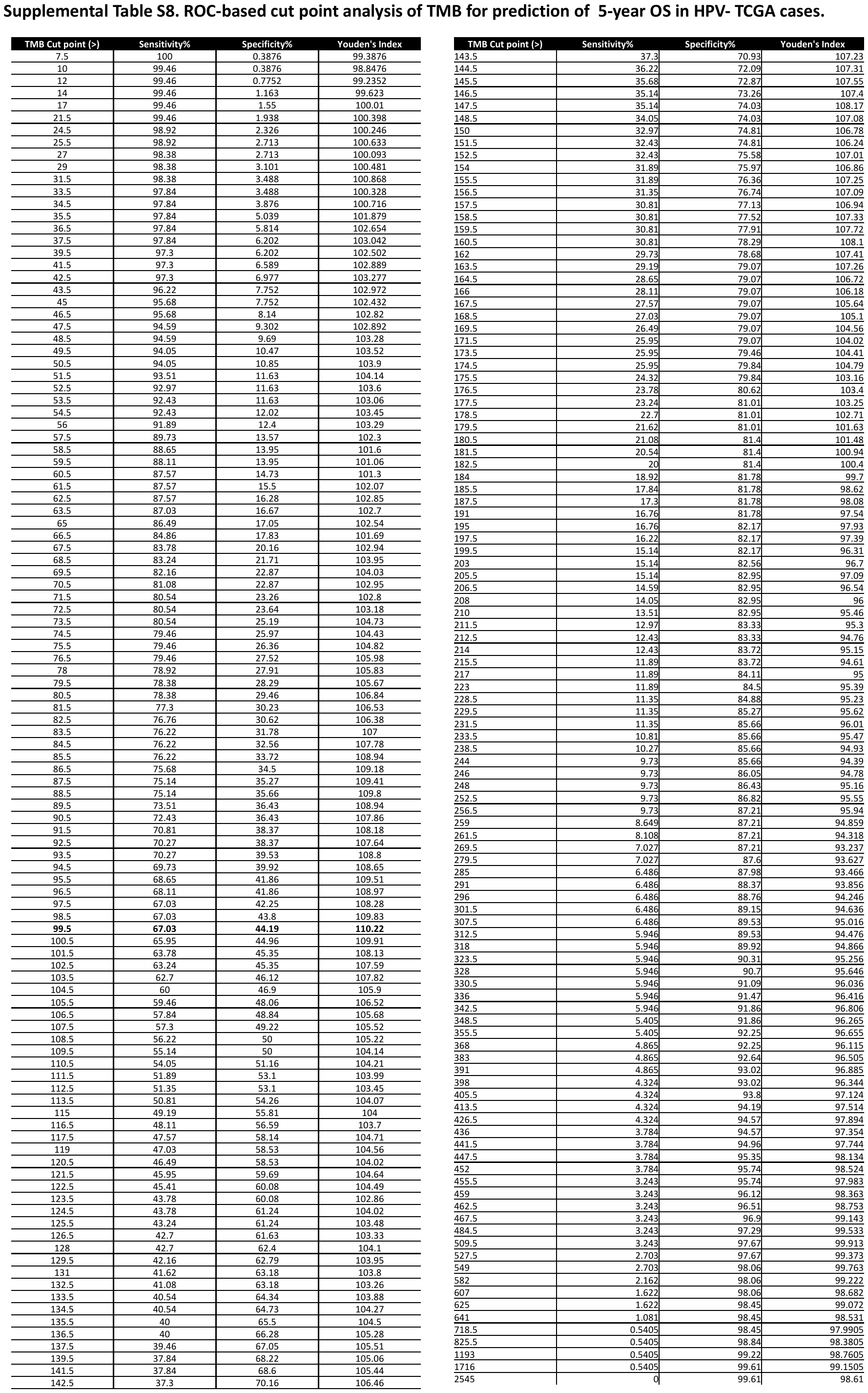

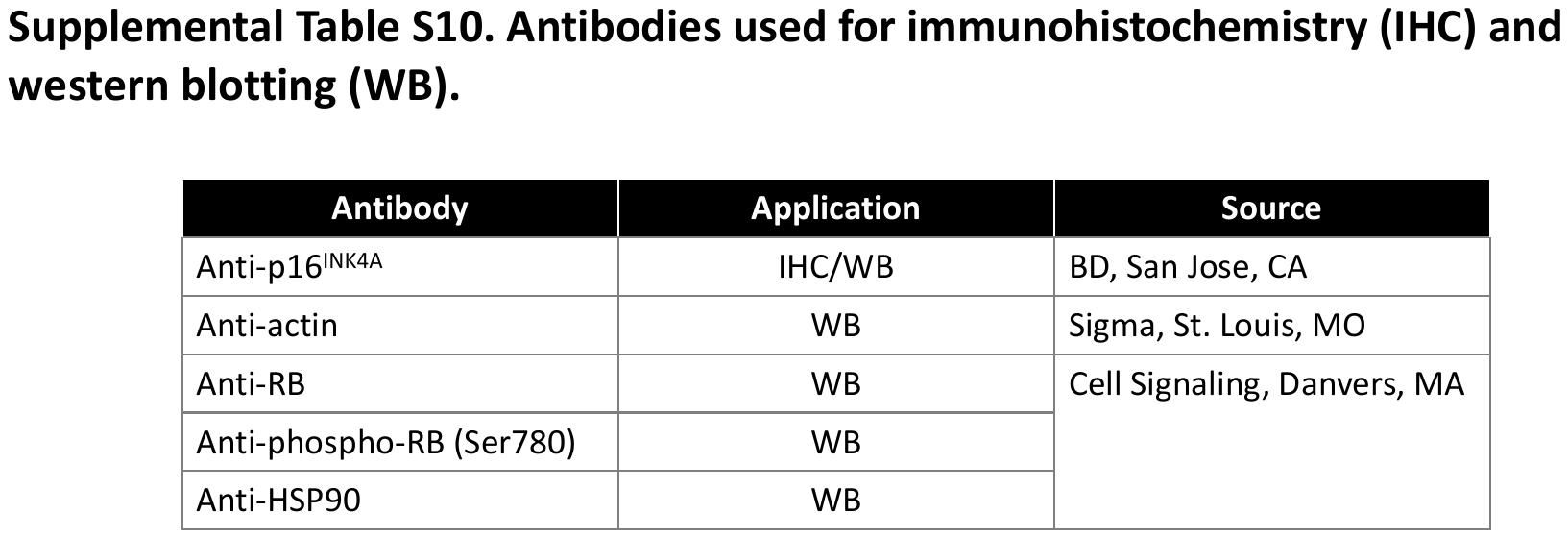
